# A Reference Data Set for Circular Dichroism Spectroscopy Comprised of Validated Intrinsically Disordered Protein Models

**DOI:** 10.1101/2023.10.19.562942

**Authors:** Gabor Nagy, Søren Vrønning Hoffman, Nykola C. Jones, Helmut Grubmüller

## Abstract

Circular Dichroism (CD) spectroscopy is an analytical technique that measures the wavelength-dependent differential absorbance of circularly polarized light, and is applicable to most biologically important macromolecules, such as proteins, nucleic acids, and carbohydrates. It serves to characterize the secondary structure composition of proteins, including intrinsically disordered proteins, by analyzing their recorded spectra. Several computational tools have been developed to interpret protein CD spectra. These methods have been calibrated and tested mostly on globular proteins with well-defined structures, mainly due to the lack of reliable reference structures for disordered proteins. It is therefore still largely unclear how accurately these computational methods can determine the secondary structure composition of disordered proteins.

Here, we provide such a required reference data set consisting of model structural ensembles and matching CD spectra for eight intrinsically disordered proteins. Using this set of data, we have assessed the accuracy of several published CD prediction and secondary structure estimation tools, including our own CD analysis package SESCA. Our results show that for most of the tested methods, their accuracy for disordered proteins is generally lower than for globular proteins. In contrast, SESCA, which was developed using globular reference proteins, but was designed to be applicable to disordered proteins as well, performs similarly well for both classes of proteins. The new reference data set for disordered proteins should allow for further improvement of all published methods.

## Introduction

Circular Dichroism (CD) spectroscopy measurements serve to estimate the average secondary structure (SS) content of proteins, to monitor protein folding under various experimental conditions, and to determine folding kinetics.^1–4^ Several CD-based SS estimation methods have been developed either as web-based applications like DichroCalc^5^, K2D3^6^, BestSel^7^, and PDB2CD^8^ or as stand-alone bioinformatics tools like SELCON3^9^, CCA^10^, and SESCA^11^. Online tools and repositories such as Dichroweb^2^ and the Protein Circular Dichroism Databank^12^ (PCDDB) also allow easy access to these tools and provide a platform for further development efforts (See Table S1 for available links).

CD spectroscopy is also often used to identify intrinsically disordered proteins (IDPs). IDPs form a major class of proteins that fulfil their biological function without adopting a well-defined secondary or tertiary structure under physiological conditions, and thus do not conform to the classical structure-function paradigm^13^. Instead, IDPs often adopt a large number of partially folded transient structures, and this conformational flexibility provides them functional advantages over their well-folded globular counterparts. Rather than forming two distinct classes, the transition between ordered and disordered proteins is continuous, and studies estimate that approximately 30% of human proteins contain flexible or disordered domains. Because of their abundance and functional importance in higher organisms, several tools have been developed to identify IDPs and intrinsically disordered regions (IDRs) in otherwise folded proteins. Most of these methods are based either on protein sequence, or the measured CD spectra of the respective regions.

The SS composition of proteins strongly affects their CD spectra. Structure-based predictions of CD spectra using quantum-mechanical calculations are challenging and computationally demanding; therefore, many CD-based analysis tools use reference data sets (RDS) instead to empirically extract structure-spectrum relationships. For folded proteins, such RDSs are available, consisting of proteins with known structures derived from X-ray crystallography, and respective CD spectra. Information from these data sets is often the basis of current algorithms that predict the CD spectrum of a putative protein structure, or infer the unknown SS composition of a protein based on its measured CD signal.

Unfortunately, the conformational flexibility of IDPs and IDRs renders them hard to characterize in terms of their structure both experimentally and computationally. Most IDPs do not form regular crystals, and if they do, e.g., in the presence of a binding partner, their crystal structure usually does not reflect their conformational flexibility in solution. Due to the lack of reliable IDP structural models, disordered proteins are largely absent from currently available RDSs, despite the fact that their CD spectra are often published, and are distinctly different from that of folded proteins.

The conformational flexibility of IDPs can be modelled through structural ensembles. Structural ensembles (or ensemble models) consist of protein conformations and associated weights that collectively describe the average protein structure and its fluctuations over time. Marked improvements in simulation force fields and molecular modelling tools now allows one to construct increasingly realistic ensemble models, which agree with or predict experimental observables. Additionally, recent developments in prediction tools can now process structural ensembles to predict observables such as, fluorescence spectra, nuclear magnetic resonance (NMR), electron paramagnetic resonance (EPR), and small angle X-ray scattering (SAXS). These developments have recently enabled more rigorous validations and further refinements of IDP ensembles.^14,15^ Additionally, online repositories such as the protein ensemble database^16^ (PED), the Biological Magnetic Resonance Databank^17^ (BMRB), and the PCDDB^12^ compile and link available data to facilitate ensemble model generation. Finally, regarding the prediction of CD spectra, our CD analysis package SESCA can predict CD spectra not only of individual protein structures, but also of structural ensembles^11^, and estimate the SS composition of proteins based on their measured CD spectrum^18^. Initially, due to the lack of available IDP RDSs, SESCA was parametrized and validated on folded proteins only.

These advances, taken together, now enable us to provide a small RDS, namely IDP8, consisting of measured CD spectra and structural ensembles for eight disordered proteins. Further, we will use this newly constructed RDS to assess the accuracy of several established modelling tools for either CD prediction or SS estimation of IDPs including the current version of our own SESCA analysis package. Our analysis indicates that the IDP8 RDS offers the opportunity not only to assess the prediction accuracy of CD-based analysis tools regarding disordered proteins, but to further improve their accuracy and precision as well.

## Materials and Methods

### Reference data set assembly

The IDP8 RDS consists of eight IDP CD spectra and 14 structural ensembles, which were assembled with the aim of testing the accuracy of CD-based prediction and SS estimation methods. The RDS includes eight disordered protein models : 1) α-synuclein (*asyn*), 2) the measles virus nucleoprotein tail domain (*mevn*), 3) *Saccharomyces cerevisiae* CDK inhibitor N-terminal targeting domain (*sic1*), 4) the human tau protein K18 fragment (*tk18)*, 5) the activator of thyroid hormone and retinoid receptor protein activation domain 1 (*actr*), 6) CREB-binding protein nuclear coactivator binding domain (*cbpn*), 7) the protein 53 N-terminal transactivation domain (*p53t*), and 8) an RS-repeat peptide (*rsp8,* sequence: GAMGPSYGRSRSRSRSRSRSRSRS). Two of these models are full length IDPs (*asyn* and *rsp8*), and the other six models (*mevn*, *sic1*, *tk18*, *actr*, *cbpn*, *p53t*) are IDRs of larger proteins. All eight disordered models were selected based on the availability of experimental and modelling data.

There are eight CD spectra included in the IDP8 RDS. We measured the CD spectra of *actr*, *asyn*, *cbpn*, *p53t*, and *rsp8* using a synchrotron radiation source (SR-CD), which allowed us to determine additional short wavelength information (down to 178 nm). The remaining three CD spectra of *mevn, sic1,* and *tk18* were measured using conventional CD spectrophotometers. Due to high absorbance and a weaker UV source in conventional CD spectrophotometers, measurements at short wavelengths are unreliable for these spectra and therefore were truncated to the wavelengths provided in Table 1. Further details are provided in the Circular Dichroism measurements section.

**Table 1:**
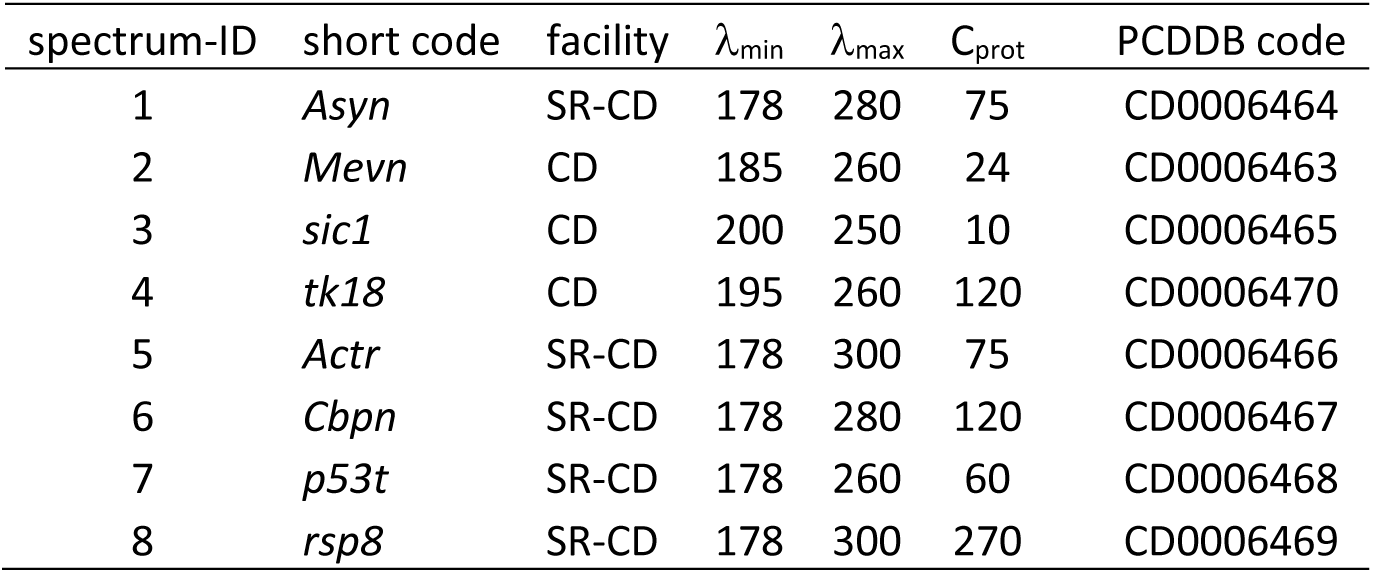
Measured IDP8 CD spectra. Properties of the eight CD spectra included within the IDP8 RDS. Columns list the ID and the short code of the protein, their minimum (λ_min_) and maximum (λ_max_) wavelengths (in nm) of the spectra, whether it was recorded on a conventional spectrophotometer (CD) or a synchrotron radiation CD (SR-CD) facility, and the estimated protein concentration (C_prot_, in µM) of the measured sample. Expected Protein Circular Dichroism Data Bank (PCDDB) accession codes for each CD spectra are also shown.

The 14 structural ensembles of the data set are organized into three groups A, B, and C based on the experimental data used in their creation. Group A contains four IDP model ensembles for *asyn*, *mevn*, *tk18*, *sic1* that were previously published on the PED^16^ under accession codes provided in Table 2. These ensembles were fitted mainly against results from NMR measurements, partly complemented by data from electron-spin paramagnetic resonance (EPR), small angle X-ray scattering (SAXS), and residual dipolar coupling (RDC) experiments. For these ensembles, CD spectra were not used during the ensemble refinement process. Group B consists of five IDP model ensembles for *mevn*, *actr*, *cbpn*, *p53t*, and *rsp8*. These ensembles were refined from large molecular dynamics (MD) simulation ensembles using the Bayesian Maximum Entropy (BME) approach to fit against measured CD spectra, SAXS curves, and NMR C_α_ chemical shifts as described below. Finally, Group C contains five model ensembles of the same five IDP domains as group B, but here, the refinement was carried out without the CD information. Separating the ensemble models into three groups allowed us to compare the average accuracy of BME refined ensemble models to established structural ensembles (group A vs. group-C), and to assess the effects of including CD spectra in the refinement process (group B vs. group C).

**Table 2:**
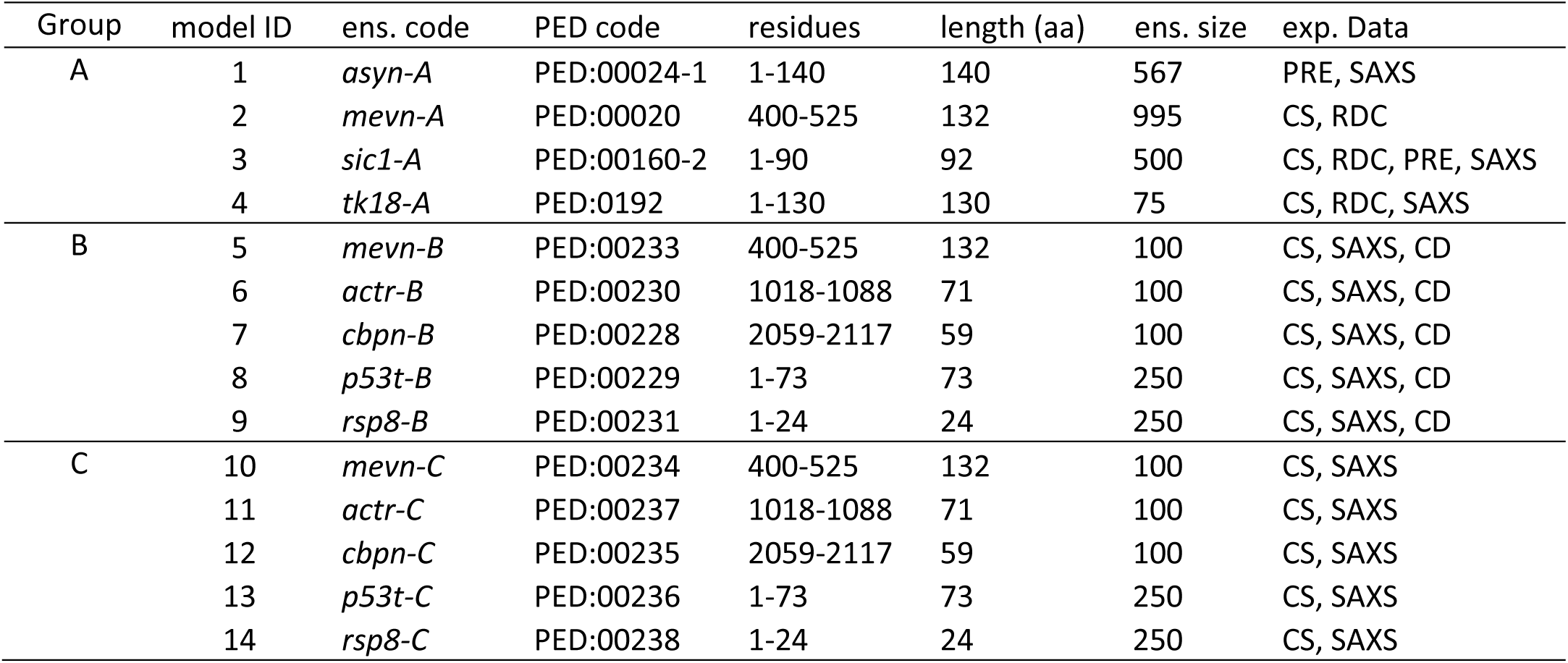
IDP8 structural ensembles. Summary of the 14 model ensembles included within the IDP8 RDS. The columns list the group and model ID as well as the short ensemble code (ens. code) of the models, the PED accession code of the model, residue numbers of the IDP domain in the full protein, the length of peptide sequence (in amino acids), the number of conformations in the model ensemble (ens. size), and experimental data used to construct or refine the model ensemble. Abbreviations of experimental data denote NMR chemical shifts (CS), NMR paramagnetic relaxation enhancement (PRE), NMR residual dipolar coupling (RDC), small angle X-ray scattering (SAXS), and Circular Dichroism (CD).

| Group | model ID | ens. code | PED code | residues | length (aa) | ens. size | exp. Data |
| --- | --- | --- | --- | --- | --- | --- | --- |
| A | 1 | *asyn-A* | PED:00024-1 | 1-140 | 140 | 567 | PRE, SAXS |
|  | 2 | *mevn-A* | PED:00020 | 400-525 | 132 | 995 | CS, RDC |
|  | 3 | *sic1-A* | PED:00160-2 | 1-90 | 92 | 500 | CS, RDC, PRE, SAXS |
|  | 4 | *tk18-A* | PED:0192 | 1-130 | 130 | 75 | CS, RDC, SAXS |
| B | 5 | *mevn-B* | PED:00233 | 400-525 | 132 | 100 | CS, SAXS, CD |
|  | 6 | *actr-B* | PED:00230 | 1018-1088 | 71 | 100 | CS, SAXS, CD |
|  | 7 | *cbpn-B* | PED:00228 | 2059-2117 | 59 | 100 | CS, SAXS, CD |
|  | 8 | *p53t-B* | PED:00229 | 1-73 | 73 | 250 | CS, SAXS, CD |
|  | 9 | *rsp8-B* | PED:00231 | 1-24 | 24 | 250 | CS, SAXS, CD |
| C | 10 | *mevn-C* | PED:00234 | 400-525 | 132 | 100 | CS, SAXS |
|  | 11 | *actr-C* | PED:00237 | 1018-1088 | 71 | 100 | CS, SAXS |
|  | 12 | *cbpn-C* | PED:00235 | 2059-2117 | 59 | 100 | CS, SAXS |
|  | 13 | *p53t-C* | PED:00236 | 1-73 | 73 | 250 | CS, SAXS |
|  | 14 | *rsp8-C* | PED:00238 | 1-24 | 24 | 250 | CS, SAXS |

### Protein sample preparation

The protein samples for four IDP domains were manufactured by the company Karebay and were delivered in a lyophilized form. The peptides were manufactured using sodium acetate buffer to avoid chloride contamination of the samples. These samples included *actr*, *cbpn*, *p53t* (13-61), and an RS repeat *rsp8*. Samples for two other variants of *p53t* (1-73 and 1-94), as well as *asyn* were kindly provided by S. Becker, Max Planck Institute for Multidisciplinary Sciences, Department of NMR-based Structural Biology, Göttingen, Germany. All seven listed protein samples were dissolved in a 10 mM sodium-phosphate buffer, pH 7.2, including 50 mM NaF for electrostatic screening. A summary of the sequence details of the IDP8 model proteins is provided in Table 2.

### Circular Dichroism measurements

Circular Dichroism (CD) spectra for seven of the protein samples described above were recorded on the AU-CD beamline of the ASTRID2 synchrotron radiation source, at the Department of Physics & Astronomy, Aarhus University, Denmark. The spectra were measured at 25 °C using a Hellma quartz suprasil cuvette type 121.000 with a nominal 0.1 mm path length under a nitrogen atmosphere. The actual path length of the cuvette was measured using an interference method^19^ to be 0.1023 ± 0.0005 mm. The CD intensities were recorded every 1 nm, with an average of 2 seconds per measurement. The final CD spectrum was calculated as the smoothed average of five independently measured and baseline corrected spectra recorded between 178-280 nm. Spectra were smoothed using a 7 pt Savitzky-Golay filter. The protein samples of *actr*, three variants of *p53t*, *cbpn*, *rsp8*, and *asyn* were measured in the buffer solution as described above. Protein concentrations for CD measurements were between 0.3 – 1.5 g/L, calculated from sample UV absorption at 214 nm.. The molar extinction coefficient at 214 nm was estimated based on the protein sequence using the method proposed by Kuipers et al with an estimated 4% uncertainty for IDPs^20^ The uncertainty of the computed concentrations was between 4% to 10% of the estimated value, based on the uncertainties of UV absorbance at 214 nm (1-5%), the pathlength (0.5%), and the extinction coefficients^20^. The protein concentration was also determined from UV absorption at 280 nm for *p53t* samples for additional validation which showed an average 19% uncertainty, and 11% average deviation from the concentrations determined from absorbance at 214 nm. Other samples lacked the sufficient absorbance at this wavelength for precise concentration determination The molar extinction coefficient at 280 nm was estimated based on Miles et al.^21^ with 12% estimated uncertainty based on the study of Pace et al.^22^ We note that due to the similarity of the measured CD spectra only one variant (1-73) of p53t was included in the IDP8 set.

The CD spectrum for *mevn* was kindly provided by Longhi et al.^23^. This spectrum was measured in a Jasco-810 dichrograph using a 1 mm quartz cuvette, 7 µM protein sample in a 10 mM sodium-phosphate buffer, pH 7.0, at 20 °C, under nitrogen atmosphere. The CD spectrum of *sic1* was kindly measured and provided by Chong et al. (personal communication). It was measured in a Jasco-1500 CD spectrophotometer using a (nominally) 0.1 mm quartz cuvette under nitrogen atmosphere, at 25°C. The measured sample had a 10 µM protein concentration, dissolved in a 50 mM potassium-phosphate buffer, pH 7, and included 150 mM NaCl and 1 mM EDTA. The CD spectrum of *tk18* was extracted from the work of Barghorn et al.^24^ In this study, the CD spectrum of tk18 was recorded using a Jasco-715 CD spectrometer, a 5 mm cuvette, and a standard PBS buffer at pH 7.4 with a protein concentration in the 50-600 µM range determined UV absorbance at 214 nm.

### Small Angle X-ray scattering

Small-angle X-ray scattering curves were measured for *actr*, three *p53t* variants, *cbpn*, and *asyn* at the European Synchrotron Radiation Facility (ESRF) Grenoble, France, at the BioSAXS beamline BM29 in 2018. All measurements were carried out under sample flow to reduce the effects of radiation damage during the measurement. SAXS curves were collected over 10 data frames of 0.3 seconds each. The measured scattering curves were normalized for the protein concentration, corrected for the buffer signal, and averaged to obtain the final scattering curves. Data processing and automated analysis was done using the Edna software package^25^. Samples were measured under similar conditions as described above, with protein concentrations ranging from 2-8 g/L.

The SAXS curve for *mevn* was kindly provided by Longhi et al., measured at the European Synchrotron Radiation Facility (ESRF) using a 10 mM Tris/Cl buffer (pH 8) containing 10% glycerol and 600 µM *mevn* at 8 °C. The SAXS curve for *rsp8* was kindly provided by Rauscher et al., which was measured at 25 °C in a 50 mM sodium-phosphate buffer (pH 7), at a concentration of 750 µM *rps8* and 100 mM NaCl. The SAXS curves for *tk18* and *sic1* were extracted from the studies of Mylonas et al. ^26^and Mittag et al. ^27^, respectively.

### Nuclear Magnetic resonance chemical shifts

Backbone chemical shifts for *asyn*, *sic1*, *tk18*, *actr*, *cbpn*, *p53t* were downloaded from the Biomagnetic Resonance database (BMRB)^17^: entry numbers 19257, 16657, 19253, 15397, 16363, and 17660, respectively. Backbone chemical shifts for *mevn* were measured by Gely et al.^28^ and kindly provided by S. Longhi. The chemical shifts were determined for a 500 µM *mevn* sample in 10 mM sodium-phosphate buffer with 50 mM NaCl, 1 mM EDTA, and 5% D_2_O, at 25 °C, pH 6.5. The backbone chemical shifts of *rsp8* were measured and kindly provided by Rauscher et al^29^. These chemical shifts were measured for a 750 µM peptide sample in a 50 mM sodium-phosphate buffer, 100 mM NaCl, at 25 °C and pH 7.

### Molecular Dynamics simulations

All-atom molecular dynamics (MD) simulations for *mevn*, *actr*, *cbpn*, *p53t*, and *rsp8* were carried out using the GROMACS 2019 software package^30^. All simulations were performed at a constant temperature of 25 °C, constant pressure of 1 atm in a dodecahedral simulation box filled with explicit water molecules and periodic boundary conditions. To accommodate extended IDP conformations, simulation box radii were chosen to be larger than the expected radius of gyration by at least 2.5 nm, resulting in system sizes of 60 000-300 000 atoms. Sodium and chloride ions were added to all simulation boxes to obtain neutral systems with NaCl concentrations of 50-150 mM. Details on simulation conditions, used force fields, and length of simulation trajectories for individual IDPs are provided in Table S2.

The temperature was kept constant by using the velocity rescaling algorithm^31^ and a coupling constant of 0.1 ps. Pressure was maintained by the Parrinello-Rahman barostat^32^ using a 0.1 ps coupling constant and the isothermal compressibility of water 4.5×10^−5^ bar^−1^. Simulations were propagated using a leapfrog integrator^33^ with 4 fs time steps. To enable such large time steps, fast vibrational degrees of freedom were removed by using the LINCS algorithm^34^ and applying a sixth order iterative restraint on the bond angles. Apolar hydrogen positions were described using virtual atom sites^30^ to eliminate hydrogen bond vibrations. Electrostatic and van der Waals interactions were explicitly calculated within a cutoff distance of 1.0 nm. Electrostatic interactions beyond the cutoff distance were calculated by particle-mesh Ewald summation ^35^ with a grid spacing of 0.12 nm. Long-range van der Waals dispersion corrections^36^ were applied to the total energy of the system in all simulations.

MD trajectories were generated using six different force fields, for which the accuracy for IDPs has been assessed previously ^29,37,38^. These force fields include the Amber03 force field^39^ with a modified TIP4P water model^40^, an Amber99SB parameter set with a modified TIP4P water model^40^, the Amber99SB-disp force field with a modified TIP4P water model re-parametrized with dispersion corrections^41^, the Amber14SB^42^ force field with an Optimal Point Charge (OPC) water model^43^, the CHARMM22* force field^44^ with a modified TIP3P water model^45^, and the CHARMM36M force field with an OPC water model.

To provide initial conformations for the BME refinement, conformations were taken at 1-100 ns intervals from 5-60 MD simulation trajectories per system amounting to total simulation times of 30-800 µs. Starting conformations for these simulations were either extended disordered structures or conformations observed in the crystalized complex structures published in protein data bank (PDB) entries 1KB6, 2L14, and 1ZQO, respectively.

### Bayesian Maximum Entropy refinement

Structural ensembles of group B and C for *mevn*, *actr*, *cbpn*, *p53t*, and *rsp8* were obtained using BME refinement. Table S3 summarizes the refinement parameters used for each IDP model. For each IDP model an initial ensemble was formed from 5,000 to 50,000 conformations obtained from the all-atom MD simulations described above. Uniform prior weights were assigned to each conformation of the initial ensembles. For each conformation, CD spectra, backbone carbon chemical shifts, and SAXS curves were computed using the SESCA (V0.96), Sparta+ (V2.6), and CRYSOL (ATSAS V2.7.2.5) analysis software packages, respectively.

All conformations of the initial ensembles were reweighted using the BME approach such that the re-weighted ensemble fits *O*_*ki*_, the *i^th^*measured observable of type *k,* as best as possible, while at the same time minimizing the loss of relative entropy *S*_rel_ = − *W*_*j*_ · log (*W*_*j*_⁄*W_j_*^0^) from redistributing the conformation weights. Here, *k* and *i* denote the type and index of the observable, while *j* is the index of the conformations. The redistributed (posterior) weights *W*_*j*_ were obtained by minimizing

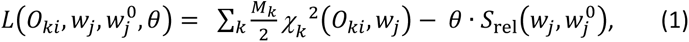

where *W_j_*^0^ are the initial (uniform) weights of each conformation, *θ* is the scaling parameter for the entropy loss, and *M*_*k*_ is the number of fitted observables of type *k*. The deviation from the observables was quantified by the *χ*^2^ deviations between each observable computed from the reweighted ensemble 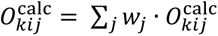 and the measured observable *O*_*ki*_

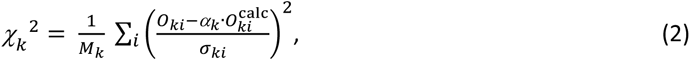

where *α*_*k*_ is the uniform scaling factor to match the measured and calculated observables of type *k*, and *σ*_*ki*_ is the uncertainty of the observable *O*_*ki*_. For group B ensembles, three types of measured observables were used for BME refinement: 1) the intensities of the measured CD spectra, 2) the SAXS intensities, and 3) the resolved C_α_ backbone chemical shifts of each residue. For group C ensembles, only SAXS intensities and C_α_ chemical shifts were used. We note that the scaling factor *α*_*k*_ was used to compensate for the machine-dependent beam intensity for SAXS measurements (*α* ∈ *R*_+_). For the CD spectrum intensities and NMR chemical shifts, no such compensation was required, hence α was set to 1.

The uncertainty *σ*_*ki*_ for SAXS measurements was defined as the SD of obtained SAXS intensities at scattering vector *q*_*i*_. The uncertainties of the backbone carbon chemical shifts *i* were set to 0.95, 1.03, and 1.13 ppm for C_α_, C_β_, and carboxylate C shifts, respectively, to reflect the uncertainty of SPARTA+ chemical shift predictions. These are conservative estimates that are considerably larger than the 0.1-0.4 ppm errors indicated in available BMRB entries of *actr*, *p53t*, and *sic1*. The uncertainty of CD intensities was computed as *σ*_*ki*_ = *δ*_*k*_ · *O*_*ki*_ + *σ_k_*^0^, where *δ*_*k*_ = 0.2 represents the typical uncertainty of intensity normalization associated with concentration and pathlength determination, and *σ_k_*^0^ = 0.75 kMRE is the machine error of CD measurements, determined from the average SD of obtained CD intensities upon repeated measurements.

The refinement parameter *θ* controls the balance between close agreement with measured observables and reducing the effective ensemble size. To find the optimal *θ* parameter for each model ensemble, several refinements with *θ* = {0.1, 1, 2, 5, 10, 20, 50, 100, 200} were carried out while monitoring the computed *χ*_*k*_^2^ values. The refinement with the largest *θ* and significant improvements to *χ*_*k*_^2^ values was selected and was used to draw sub-ensembles that constitute the final ensemble models.

To obtain the final ensemble models, smaller sub-ensembles of 5, 10, 20, 50, 100, and 200 conformations were drawn at random by rejection sampling based on the redistributed weights of the conformations after refinement. Conformations with high redistributed weights in the initial ensemble may be included multiple times in the final ensemble to represent their importance. To assess the effect of the sub-ensemble size on the model accuracy, five sub-ensembles were drawn and the deviation from experimental observables was computed and averaged for each size. The sub-ensembles of each size were concatenated to form a combined ensemble model for each IDP. Deviations from the measured observables were calculated for the concatenated ensemble as well. Finally, the ensemble with the smallest size was selected for each IDP that met the two following criteria: 1) increasing the ensemble size further does not improve the average *χ_κ_*^2^ deviations considerably, and 2) the average *χ_κ_*^2^ deviations of sub-ensembles are similar to *χ_κ_*^2^ deviations of the concatenated ensemble within uncertainty. The selected ensemble sizes and *θ* parameters for all derived IDP models are summarized in Table S3. This procedure yielded small ensemble models of 100-250 conformations with integer weights for each refined ensemble that minimize the chance of overfitting to experimental information.

### Accuracy of the predicted CD spectra

To assess the accuracy of the CD spectrum predicted for protein *j*, the predicted CD spectrum was compared to the measured spectrum by computing the root mean squared deviation (RMSD) of CD intensities,

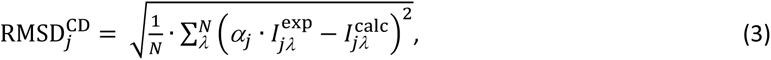

expressed in 1000 mean residue ellipticity units (kMRE, 1000 deg cm^2^/dmol) for each wavelength λ for which both the measured and the predicted spectrum were available. Here, *N* is the number of available wavelengths, and 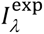 and 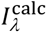 are the measured and predicted CD intensities, respectively. The scaling factor *α*_*j*_ minimizes the RMSD described above and accounts for experimental spectrum normalization errors.

To assess the accuracy of CD spectrum prediction methods for disordered proteins, the CD spectra of all model IDPs in the IDP8 RDS were predicted from their structural ensembles, the deviation from their measured reference CD spectra were computed, and the resulting RMSD^CD^ values were averaged to determine the mean accuracy of the respective method. A similar approach was followed to assess the mean accuracy of each studied CD prediction method for globular proteins, using the reference structures and CD spectra of the SP175 RDS.

### Accuracy of estimated SS fractions

To determine the accuracy of SS estimation methods, the RMSD of SS fractions was computed for each protein *j* in globular and disordered protein RDSs as follows. For globular proteins of the SP175 RDS the SS fractions estimated from their CD spectra were compared to the respective reference structures derived from X-ray diffraction measurements. For the disordered proteins of the IDP8 RDS, estimated SS fractions were compared to those computed from the reference ensemble models. The RMSD between the estimated and reference SS was computed as

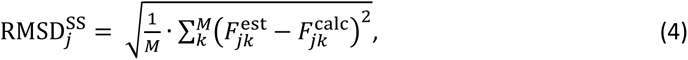

where M is the number of SS classes within the classification method, and 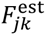 and 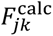 are the estimated and computed fractions of SS class *k*, respectively.

To be able to apply the RMSD determination according to eq. 4, the SS fractions computed from the reference structures/ensembles by an SS classification method have to be grouped and identified with the classes of the SS estimation method. For SESCA, the documented calculation of SS fractions from the protein structure was used. For basis sets DS-dTSC3 and DS5-4SC1, the SS composition was computed using the DISICL algorithm, and the SS elements were grouped into three and six SS classes, respectively. For the DSSP-1SC3 basis set, the SS composition was determined using the DSSP algorithm and the obtained SS fractions were grouped into four SS classes. For the HBSS-3SC1 basis set, the HBSS algorithm was used, and the obtained SS composition was grouped into five SS classes. To assess the accuracy of the K2D3 algorithm, SS classification was performed the same way as for the DS-dTSC3 SESCA basis set, and the Alpha(-helix) and Beta(-sheet) sheet fractions were compared to the corresponding estimated SS contents. The third SS fraction (Coil) for the K2D3 composition was computed as *F*_*j*,*Coil*_ = 1 − (*F*_*j*,*Beta*_ + *F*_*j*,*Alpha*_).

Finally, to assess the accuracy of the BESTSEL SS estimates, the SS fractions of the reference models were determined by the HBSS algorithm, which uses similar helix and advanced β-sheet classifications. The obtained fractions were grouped into six SS classes as follows: The Helix-1 and Helix-2 classes of BESTSEL were grouped into a common Helix class, which was identified with the 4-Helix class in HBSS. The three anti-parallel β-sheet classes (Anti1-3) were kept separate and were identified with the corresponding HBSS classes (left-handed, non-twisted, and right-handed β-strands). All parallel β-strand classes in HBSS were merged and identified with the parallel β-sheet class of BESTSEL. The SS fractions of all other classes in BESTSEL and HBSS were merged and identified with an “Other” SS class, resulting in six SS classes for both algorithms.

## Results and discussion

### Model quality assessment

First, we assessed how well the models of the IDP8 RDS shown in Fig. 1 agree with SAXS and NMR chemical shift measurements (agreement with CD spectra will be discussed below). Table 3 shows how well the observables predicted from the model ensembles of the RDS agree with the measured SAXS data as well as with Cα, Cβ and carbonyl-carbon (CO) chemical shifts. These two groups of observables were chosen due to their complementarity; whereas SAXS curves report overall IDP compactness, carbon chemical shifts are sensitive to the local secondary structure.

**Figure 1:**
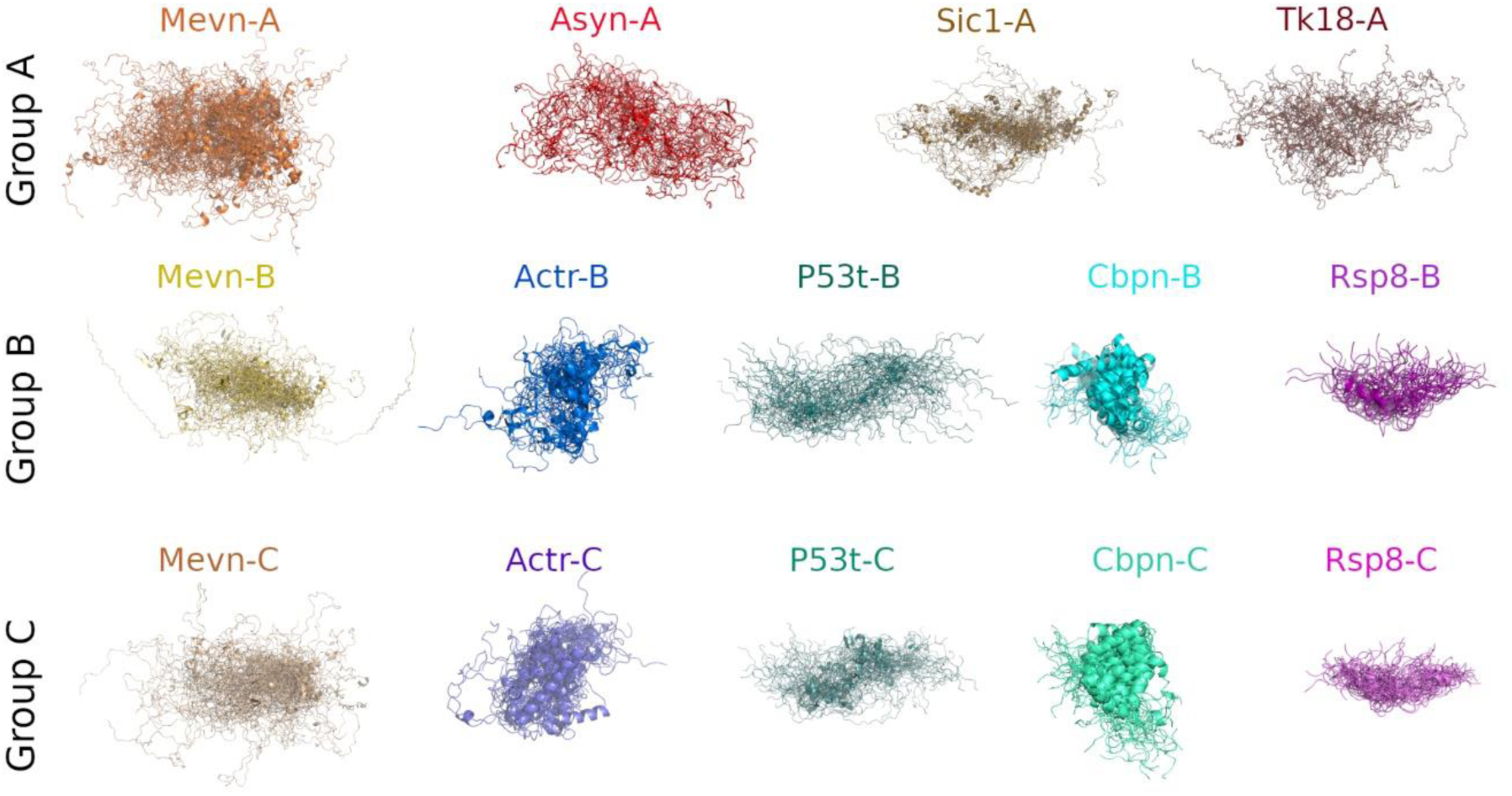
IDP8 protein ensemble models. Each ensemble model is an overlay of 20-50 backbone conformations, shown in cartoon representation, and fitted to the first model of the respective ensemble. The name of each ensemble model is displayed above the model. Group A models were previously published and were obtained from the PED, group B models were derived by the authors using NMR chemical shifts, SAXS, and CD measurements. Group C models were derived similarly as the models of group B but without using CD information.

**Table 3:** IDP8 ensemble model assessment. Summary of IDP8 ensemble model prediction vs. measured SAXS curves, NMR chemical shifts, and CD spectra. The table lists the group ID, ensemble ID, the square root of the χ^2^ deviation of the predicted and measured SAXS curves, the average RMSDs between backbone NMR chemical shifts (CS) for Cα, Cβ, and carbonyl carbon (CO) atoms, as well as the RMSD of CD intensities (CD) as predicted using the SESCA basis set DS-dTSC3. Group A comprises four reference ensembles that were available in the PED prior to this work, generated using SAXS and NMR data, but not CD spectra. Group B and C comprises ensembles that were selected from large initial ensembles generated by molecular simulations using BME refinement with SAXS and Cα chemical shift data. For the ensembles of group B, the refinement also included CD information. Group 0 comprises of the five initial ensembles group B and C were refined from. They are not part of the IDP8 data set and their deviations from the measured observables are shown here to demonstrate the effects of ensemble refinement.

| Group | ensemble code | SAXS $\chi$ | CS-Cα ppm | CS-Cβ ppm | CS-CO ppm | CD kMRE |
| --- | --- | --- | --- | --- | --- | --- |
| A | *asyn-A* | 1.34 | 0.36 | 0.66 | 0.66 | 1.9 |
|  | *mevn-A* | 0.61 | 0.28 | 0.37 | 0.40 | 2.0 |
|  | *sic1-A* | 0.66 | 1.30 | 0.49 | NA | 3.1 |
|  | *tk18-A* | 0.96 | 0.56 | 1.01 | 0.52 | 1.4 |
| B | *mevn-B* | 0.39 | 0.34 | 0.38 | 0.52 | 1.2 |
|  | *actr-B* | 1.94 | 0.35 | 0.35 | 0.46 | 0.5 |
|  | *cbpn-B* | 1.65 | 0.69 | 0.38 | 0.62 | 1.6 |
|  | *p53t-B* | 0.92 | 0.36 | 0.34 | 0.53 | 1.3 |
|  | *rsp8-B* | 1.03 | 0.41 | NA | 0.60 | 1.0 |
| C | *mevn-C* | 0.37 | 0.34 | 0.32 | 0.37 | 1.4 |
|  | *actr-C* | 1.83 | 0.28 | 0.32 | 0.39 | 3.6 |
|  | *cbpn-C* | 1.64 | 0.67 | 0.40 | 0.62 | 2.0 |
|  | *p53t-C* | 0.91 | 0.28 | 0.32 | 0.47 | 2.7 |
|  | *rsp8-C* | 1.07 | 0.57 | NA | 0.64 | 2.5 |
| 0 | *mevn-0* | 0.71 | 0.53 | 0.35 | 0.64 | 2.0 |
|  | *actr-0* | 1.72 | 0.50 | 0.36 | 0.53 | 3.4 |
|  | *cbpn-0* | 2.94 | 0.93 | 0.49 | 0.70 | 2.5 |
|  | *p53t-0* | 0.92 | 0.39 | 0.29 | 0.72 | 2.1 |
|  | *rsp8-0* | 1.67 | 0.83 | NA | 0.59 | 2.9 |

The χ values for SAXS curves shown in the second column of Table 3 are square roots of the *χ*^2^ metric defined by Sevrgun et al. This metric is insensitive to any scaling differences between the measured and predicted SAXS intensities, and reports the deviation in units of the experimental uncertainty determined by *σ*_*i*_, the standard deviation of the scattering intensities. In our measurements, this experimental uncertainty in the 0.01 to 0.25 Å^−1^ range where the radius of gyration information is typically determined varies between 2-20% (with an average of 12%) of the absolute scattering intensity. Therefore, χ values between 1.0 and 2.0 correspond to average 12-24% relative deviations from measured SAXS intensities. For seven of the 14 ensemble models, the χ values are below one, meaning that predicted SAXS intensities are on average well within the experimental uncertainty. The remaining seven models achieved χ values between one and two, resulting in an overall average χ of 1.14 for the whole RDS. This result suggests that the size distributions of the model ensembles agree with the available experimental data, except for the *actr* and *cbpn* ensembles for which the predicted SAXS curves deviate from the experiment with χ values between 1.8 and 2.0.

The *actr* and *cbpn* ensembles showcase that the BME refinement avoids overfitting to the experimental data. Briefly, BME refinement reweights conformations of an initial ensemble to achieve good agreement with experimental data (within 2 σ*_i_*), but penalizes severe deviations from the initial weights. For *cbpn* the initial MD-based ensemble prior to refinement (*cbpn-0)* had a poor agreement with SAXS data (χ of 2.94), likely due to under-sampling moderately extended helical conformations. The BME refinement increased the weights of these under-sampled conformations, consequently, both *cbpn-B* and *cbpn-C* agree significantly better (just within 2 *σ*_*i*_) with the measured SAXS data. For *actr-0* the initial deviation of from SAXS data was already acceptable within the experimental uncertainty (χ of 1.72), and increased slightly during the refinement for both *actr-B* (1.94) and *actr-C* (1.83) to improve the agreement with NMR chemical shifts.

Columns three to five in Table 3 report the root-mean-squared deviation (RMSD) of carbon chemical shifts for each model ensemble. The average RMSDs of the data set are 0.46 ppm, 0.49 ppm, and 0.46 ppm for Cα, Cβ, and CO chemical shifts, respectively. These RMSD values are slightly larger than the 0.1 - 0.4 ppm estimated experimental uncertainty reported in BMRB entries, but are considerably smaller than the average 1.14 ppm, 0.94 ppm, and 1.09 ppm backbone chemical shift deviations reported by Shen and Bax obtained in the context of Sparta+ prediction assessments from high-quality crystallographic structures of globular proteins. Considering that the local secondary structure causes approximately 7 ppm variation in carbon chemical shifts, the observed deviations would constitute 5-10% of this variation for IDP chemical shifts and 12-17% for those of globular proteins.

To assess the effects of using CD spectrum information in ensemble refinement, we compared the average deviations between predicted and measured SAXS and NMR data for ensembles of group A, B, and C separately. In addition, we also computed the SAXS and NMR deviations of the initial MD ensembles (henceforth group 0) group B and C ensembles were refined from. SAXS intensities and Cα chemical shifts were used as fit variables during both group B and group C ensemble refinements.

The average deviation from measured SAXS curves is within the average uncertainty for group A, as shown by a mean χ value of 0.87 ± 0.17. Refinement reduced the deviation from measured SAXS curves from an initial χ of 1.59 ± 0.39 for group 0 to 1.19 ± 0.27 for group B and 1.17 ± 0.26 for group C, showing no significant difference between the two groups. The average deviation of Cα chemical shifts for group A is 0.62± 0.2 ppm, which is very similar to the deviation of 0.63 ± 0.1 ppm for initial group 0 ensembles. Ensemble refinement improved the average deviation from measured Cα chemical shifts to 0.43 ± 0.1 ppm for both group B and group C ensembles.

The deviations from measured Cβ and CO chemical shifts were not used in ensemble refinements, and thus are used for cross-validation. The average deviation of Cβ chemical shifts in group A is 0.63 ppm. In comparison, the Cβ chemical shifts are accurately reproduced by the initial MD ensembles with an average Cβ shift deviation of 0.37 ± 0.04 ppm. Apparently, the refinement process did not cause significant changes in the Cβ chemical shift deviations for group B or C within the uncertainty. In contrast, average deviations from measured CO chemical shifts improved from an initial value of 0.64 ± 0.03 ppm in group 0 to 0.55±0.03 ppm for group B ensembles, and to 0.50±0.06 ppm for group C ensembles, which are small but statistically significant improvements. The ensembles in group A are similarly accurate in predicting CO chemical shifts with an average deviation of 0.53±0.06 ppm.

In summary, our structural ensembles reproduced both the measured SAXS curves and NMR chemical shifts for all model IDPs with deviations from the measurements close to the experimental uncertainty. The average agreement with SAXS curves and NMR chemical shifts indicates that there are only minor differences between the quality of published PED models in group A and the newly refined ensemble models of groups B and C. The ensembles of groups B and C also showed no significant accuracy difference regarding the predicted SAXS curves and NMR chemical shifts, suggesting that they are of similar quality. Further, comparison to the initial group 0 ensembles indicate that ensemble refinement did improve the of agreement with experimental data significantly, but these improvements did not happen at the cost of overfitting to the experimental observables. Based on the presented quality assessment, we consider the model ensembles sufficiently accurate that they can now be used to assess the accuracy of both structure-based CD prediction methods as well as CD-based SS estimation methods regarding IDPs.

### Testing CD prediction methods

Utilizing the new IDP8 RDS, we proceed to determine the accuracy of the three structure-based CD-spectrum prediction methods SESCA, PDB2CD, and DichroCalc, and compare their mean accuracy separately for IDPs and globular proteins. Figure 2 shows the eight measured CD spectra of the IDP8. The predicted CD spectra of all methods for the IDP8 RDS are compared with the measured CD spectra in Figures S1-S6. The accuracy of these algorithms on globular proteins was previously assessed using the SP175 RDS, which contains 71 water soluble globular proteins. The same SP175 data set was used as a training set for the two empirical methods SESCA and PDB2CD, with no IDPs involved. The individual RMSDs computed between the measured CD spectra of IDP8 RDS and the CD spectra predicted from the 14 ensemble models of the RDS are shown in Table 4.

**Figure 2:**
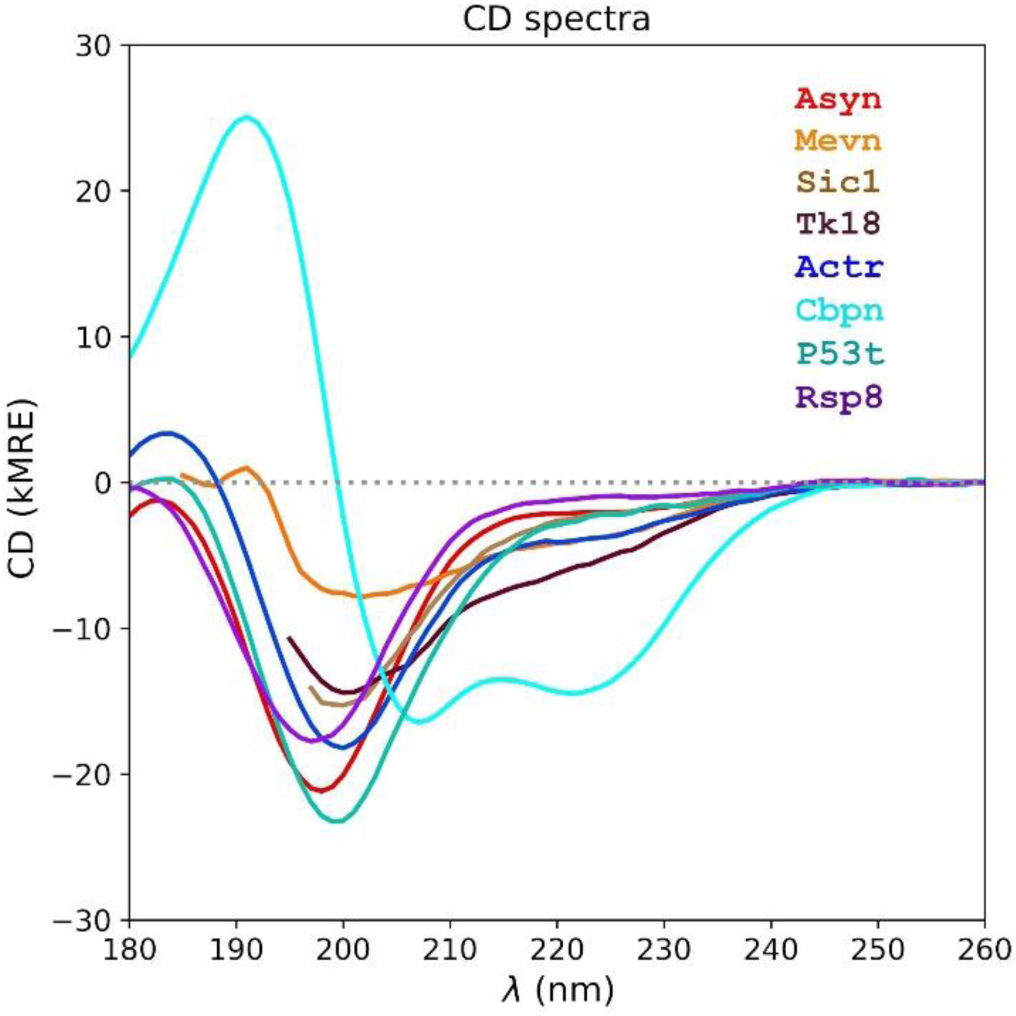
Measured IDP8 CD spectra. The spectra of eight different IDP domains are shown in different colors. Abbreviations for the name of each domain are shown in the upper right corner (color coded) and are listed in Table 1. The full name of each IDP domain is listed in the Reference data set assembly section of this manuscript. Intensities of the CD spectra are expressed in 1000 mean residue ellipticity units (kMRE or 1000 deg* cm^2^ / dmol). The dotted gray line indicates the CD intensity of 0 kMRE.

**Table 4:** Accuracy of CD spectrum predictions. Summary of RMSDs between measured CD spectra, and CD spectra predicted from IDP8 reference ensemble models. The RMSD of CD predictions using four SESCA basis sets (DS-dTSC3, DSSP-1SC3, HBSS-3SC1, DS5-4SC1), and methods PDBMD2CD and DichroCalc (Dichro) are shown in separate columns for each ensemble. RMSD values are expressed in 1000 Mean Residue Ellipticity (kMRE) units. The most accurate predictions are indicated in bold, and RMSD values for the basis set used in the ensemble refinement of group B are underlined. The average (avg) and standard deviations (SD) of the RMSD values for each basis set are shown at the bottom of the table.

| Group | ens. code | DS-dTSC3 | DSSP-1SC3 | HBSS-3SC1 | DS5-4SC1 | PDBMD2CD | Dichro |
| --- | --- | --- | --- | --- | --- | --- | --- |
| A | asyn-A | 1.92 | **1.91** | 1.55 | 1.92 | 6.31 | 8.48 |
|  | mevn-A | 1.95 | **1.87** | 1.91 | 2.88 | 3.63 | 5.50 |
|  | sic1-A | 2.67 | 2.39 | **0.57** | 2.69 | 3.60 | 3.09 |
|  | tk18-A | **1.36** | 1.40 | 1.55 | 3.18 | 2.74 | 11.59 |
| B | mevn-B | <u>1.22</u> | **1.17** | 1.60 | 1.61 | 4.74 | 8.76 |
|  | actr-B | 2.79 | 2.60 | 2.66 | **0.45** | 6.26 | 8.33 |
|  | cbpn-B | 1.07 | 1.60 | **1.06** | <u>1.47</u> | 2.87 | 5.42 |
|  | p53t-B | **1.33** | 1.43 | 1.66 | 1.89 | 6.41 | 15.61 |
|  | rsp8-B | 2.74 | 2.07 | 4.29 | **0.96** | 6.91 | 10.49 |
| C | mevn-C | 1.41 | **1.20** | 1.66 | 2.73 | 4.61 | 9.86 |
|  | actr-C | 4.89 | 3.95 | 3.97 | 3.59 | 7.02 | 6.97 |
|  | cbpn-C | 1.09 | 1.85 | **1.06** | 1.21 | 3.23 | 5.08 |
|  | p53t-C | 2.66 | 1.35 | 2.25 | **0.64** | 6.98 | 13.41 |
|  | rsp8-C | 3.07 | **1.83** | 4.34 | 2.48 | 7.08 | 9.33 |
| avg |  | 2.2 | 1.9 | 2.2 | 2.0 | 5.2 | 8.7 |
| sd |  | 1.1 | 0.7 | 1.2 | 1.0 | 1.7 | 3.4 |

Figure 3 shows the average RMSD values between measured and predicted CD spectra (RMSD^CD^, see eq 3.) for both disordered (IDP8, blue) and globular (SP175, orange) proteins. For SESCA predictions, four different basis sets were used: DS-dTSC3, DSSP-1SC3, HBSS-3SC1, DS5-SC1. These basis sets represent ‘pure’ CD spectra for given SS elements (α-helix, β-sheet etc., see Nagy et al. ^11^ for precise definitions), and therefore differ depending on which and how many SS elements have been used, as well as on which SS classification method (e.g., DISICL^5–8,11,18,46–50^, DSSP, or HBSS) has been applied. All four chosen basis sets contain correction terms for side chain signals for improved accuracy. In addition, SESCA applies intensity scaling that minimizes RMSD^CD^ values to account for potential normalization errors of the measured CD intensities. These errors are usually caused by uncertainties of the intensity calibration, protein concentration and cuvette pathlength determination. To provide a fair comparison, we also applied intensity scaling when determining the RMSD^CD^ values for PDBMD2CD and Dichrocalc predictions as well.

**Figure 3.**
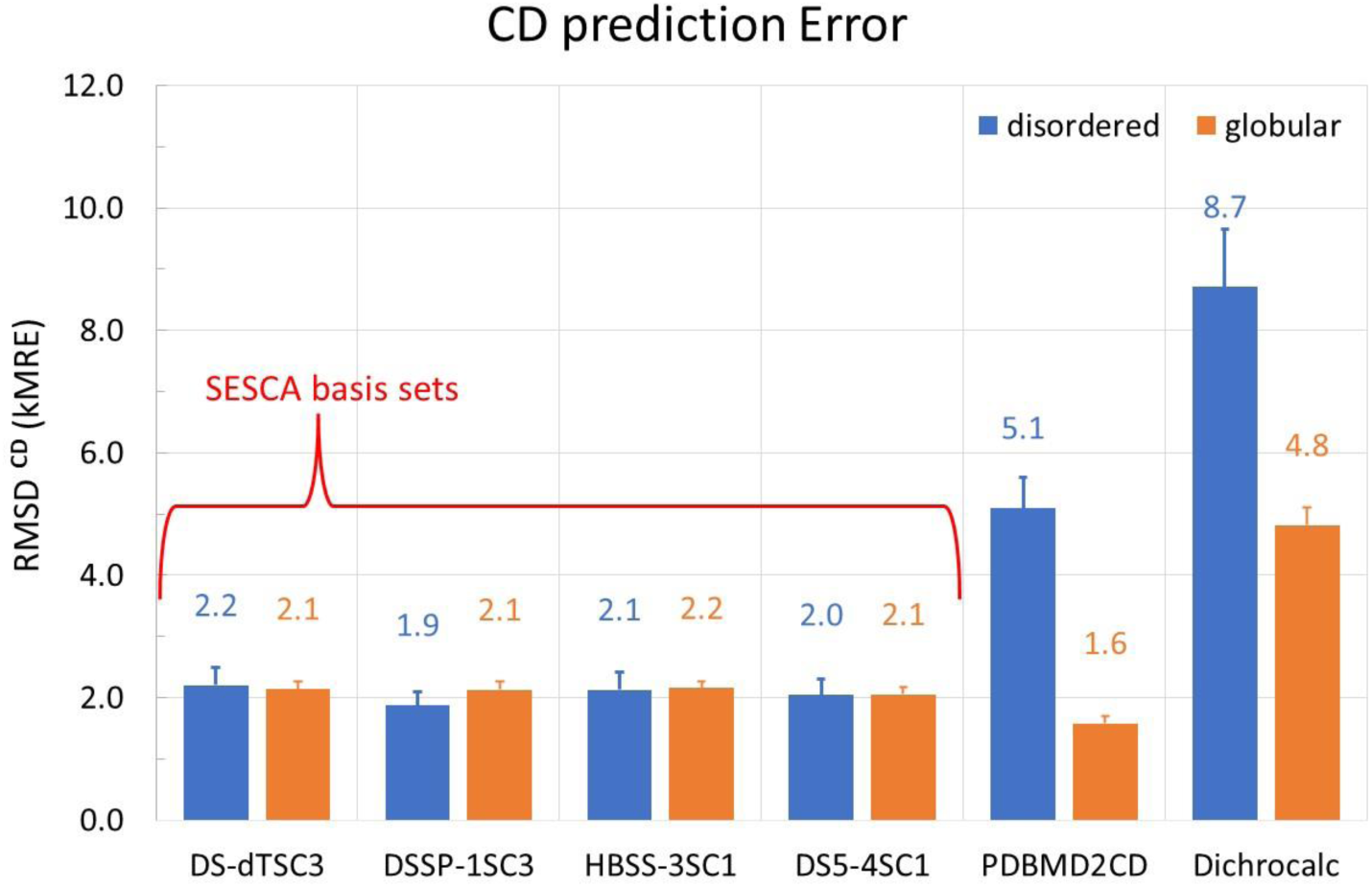
: Accuracy of CD spectrum predictions. Summary of RMSDs of CD spectra predicted from reference model structures relative to measured spectra of the same protein. Shown are RMSD values averaged over all proteins, for the different methods described in the text. Two reference data sets have been used: IDP8 for disordered proteins (blue) and SP175 for folded globular proteins (orange). Tested CD prediction methods are: DichroCalc, PDBMD2CD, and SESCA with four different basis sets (DS-dTSC3, DSSP-1SC3, HBSS-3SC1 and DS5-4SC1).

The average prediction accuracy of SESCA is 2.0 ± 0.1 kMRE for disordered proteins. As shown in Fig. 3, the average accuracy is similar for all four chosen basis sets, ranging between 1.9 and 2.2 kMRE with a mean standard deviation (SD) of 1.0 kMRE for RMSD^CD^ values within the IDP8 RDS using the same basis set. The average scatter of RMSD^CD^ values is 0.67 kMRE, when the measured CD spectra are compared to CD predictions from the same ensemble model using different basis sets. In comparison, the average prediction accuracy of SESCA for globular proteins is 2.1 ± 0.05 kMRE units (as determined from the SP175 RDS). The scatter of RMSD^CD^ values for globular proteins is 1.0 kMRE within the RDS using the same basis set and 0.7 kMRE between predictions from the same crystal structure using different basis sets. The obtained RMSD values do not show a significant difference in prediction accuracy between the chosen basis sets. Most importantly, the RMSD^CD^ values support our previous expectations that, by construction, SESCA should yield a similar accuracy for disordered proteins as for globular proteins.

Next, we tested the accuracy of the PDB2CD algorithm and its recent update PDBMD2CD that allows CD predictions from small structural ensembles. PDB2CD is based on determining the SS composition from the model structure (or ensemble) by the DSSP algorithm and produces predicted spectra by taking a weighted sum of spectra from structurally similar reference proteins. At the time of writing, PDB2CD can utilize two globular RDS: SP175 and SMP180 to predict the CD spectra of protein models. SMP180 includes all SP175 proteins and 11 additional membrane proteins, but neither RDS includes any disordered proteins, which suggests limited accuracy for this class of proteins. PDBMD2CD is based solely on the SMP180. Therefore, we used this RDS for computing CD predictions of both globular and disordered protein spectra in our evaluation (the average accuracy for globular proteins was still determined from the RMSD^CD^ values of SP175 proteins). As can be seen in Fig. 3, the accuracy of PDBMD2CD for globular proteins is slightly better than that of SESCA, with an RMSD^CD^ of 1.6 ± 0.1 kMRE (SD 1.0 kMRE). For disordered proteins, however, the prediction accuracy of PDB2CD is markedly reduced, with an average RMSD^CD^ 5.2 ± 0.5 kMRE (SD 1.7 kMRE).

In contrast to the other two empirical algorithms, DichroCalc predictions are calculated directly from the three-dimensional protein structure through parameters derived from time-dependent quantum mechanics (QM) calculations. The obtained average prediction RMSD^CD^ values for DichroCalc are 4.8 ± 0.3 kMRE (SD 2.4 kMRE) for globular proteins, and are even larger (8.7 ± 1.0 kMRE, SD 3.4 kMRE) for disordered proteins. The obtained deviations from measured CD spectra indicate that the approximations that allow CD calculations for entire proteins are rather harsh and limit the accuracy of DichroCalc in reproducing the fine spectral features. These limitations are particularly severe for disordered proteins, because the negative peak that defines the shape of their spectra is not reproduced well by the underlying matrix method.

Further, to assess the effect of using CD information during ensemble refinement we also compared the average accuracy of CD predictions of group B ensembles with those of group C ensembles shown in Table 4. Here, we will focus on the prediction accuracies of SESCA, because the large mean and scatter of RMSD^CD^ values for PDBMD2CD and DichroCalc renders it difficult to infer statistically relevant statements about model quality using these methods. During the refinement of group B ensembles, the SESCA basis set DS-dTSC3 was used to compute the CD signal of individual conformations for *mevn-B* and *p53t-B*, whereas the DS5-4SC1 basis set was used for *actr-B*, *cbpn-B*, and *rsp8-B*. The individual RMSD^CD^ values (underlined in Table 4) for CD predictions using these ensembles and the corresponding basis sets average to 1.1 ± 0.2 kMRE. This accuracy can be considered the best accuracy achievable by using our BME refinement framework, which allows us to modify the ensemble populations to better match the measured CD spectra without overfitting the experimental data. It is also a considerable improvement over the 2.6 ± 0.3 kMRE average CD deviation of the initial MD ensembles (see Table 3). The average deviation of group B ensemble CD predictions using all four chosen SESCA basis sets (lines 5-9 in Table 4) amounted to 1.8 ± 0.2 kMRE. In comparison, the average CD deviation for group C ensembles (lines 10-14) is 2.4 ± 0.4 kMRE, which suggests that including CD data in the ensemble refinement process reduces both the mean and the scatter of RMSD^CD^ values to a small but statistically significant extent.

In summary, based on the CD predictions for our IDP8 RDS, SESCA consistently predicts the CD spectra of IDPs with an accuracy similar to that of globular proteins. Additionally, SESCA predictions are robust with respect to the choice of basis set both for folded proteins and IDPs. In contrast, PDBMD2CD and DichroCalc predictions are markedly less accurate regarding IDPs than for the folded proteins. Based on our model quality assessments, including CD information during the ensemble refinement process significantly improves CD predictions from the ensemble models, while maintaining the accuracy of predicted SAXS curves and carbon chemical shifts.

### Testing IDP SS estimation methods

Next, we focused on SS estimation, the second main branch of CD-based methods, which infers the average SS composition of proteins from their measured CD spectra. Here, we assessed the SS estimation accuracy of the Bayesian SS estimator SESCA_bayes, using the same four basis sets as above, as well as two other widely used methods, namely BeStSel and K2D3. The estimated SS fractions of all methods for the IDP8 RDS are shown in Tables S4-S9. To assess the accuracy of estimated SS compositions, we compared them to reference SS compositions (see Methods Section for details). For globular proteins, SS compositions of the NMR/crystallographic structures of the SP175 RDS were used as reference. For disordered proteins, we selected the SS composition of those ensemble models from IDP8 as reference that had the lowest average RMSD^CD^ for SESCA predictions, namely *asyn-A*, *mevn-B*, *sic1-A*, *tk18-A*, *actr-B*, *cbpn-B*, *p53t-B*, and *rsp8-B*. The accuracy of the estimated SS content was quantified by the RMSD to the reference SS fractions (RMSD^SS^, see *eq.* 4). The summary of all RMSD^SS^ values shown in Table S10 indicates, that the choice of reference ensemble (except for *mevn*) doesn’t have a large impact on the average accuracy of SS estimation methods and wouldn’t change our conclusions outlined below. For *mevn* all three tested methods estimated SS fractions in better agreement with the *mevn-B* ensemble than *mevn-A or mevn-C*.

Figure 4 compares the average SS estimation accuracies of these methods for the IDP8 RDS (in blue) of disordered proteins with those obtained for globular proteins of the SP175 RDS (orange). Overall, the tested methods performed more similarly to one another than the CD prediction methods, albeit larger differences are seen between the four SESCA basis set variants. All methods achieved average RMSD^SS^ values between 0.07 and 0.12 for globular proteins and slightly larger average RMSD^SS^ values (between 0.07 and 0.14) for disordered proteins. No clear correlation is observed between the SS estimation accuracy and the number of SS classes used for the estimation method, although the precision of SESCA_bayes estimates increased monotonically with the number of SS classes in the basis set.

**Figure 4:**
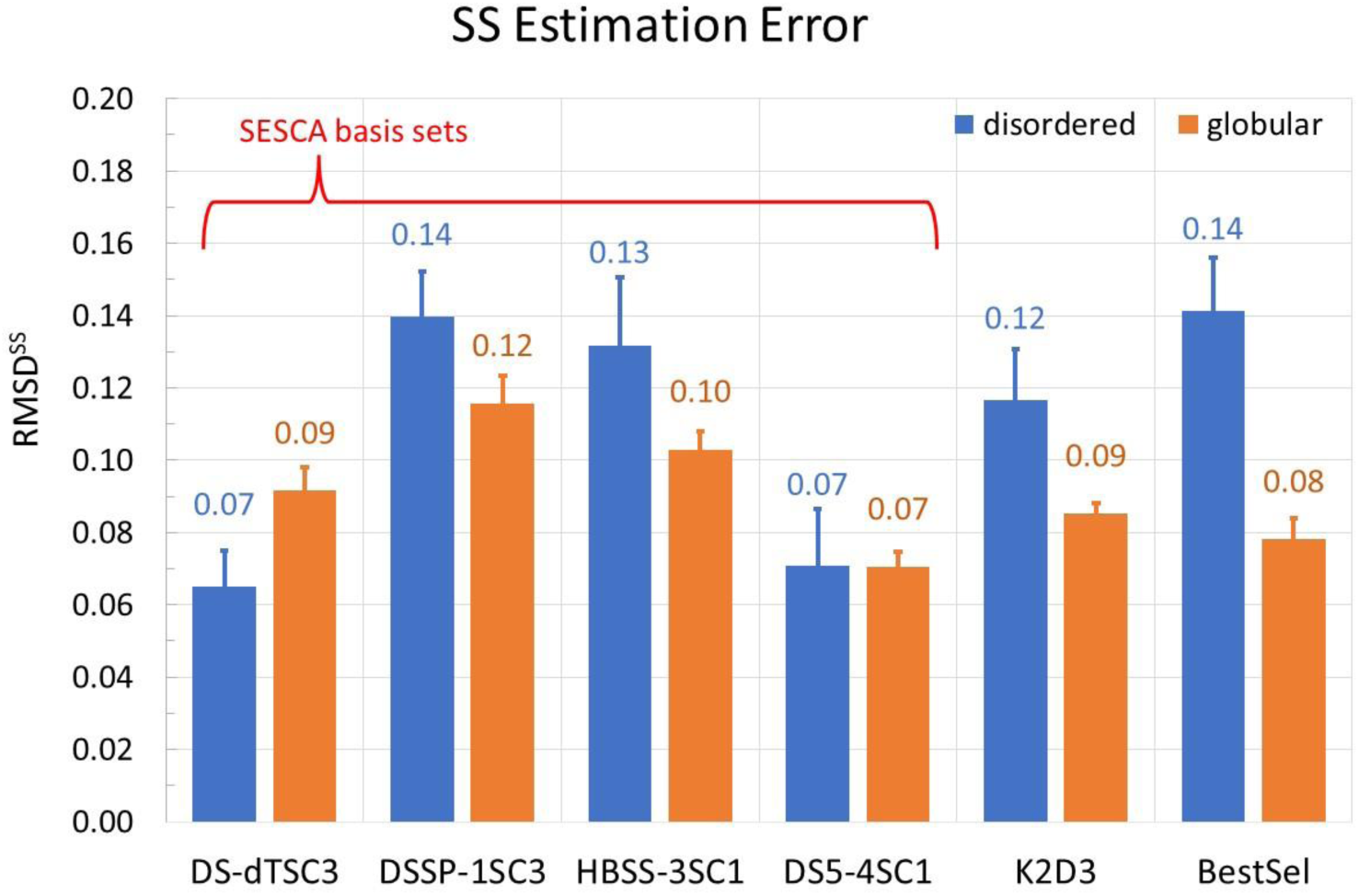
Accuracy of SS fraction estimates. Summary of averaged RMSDs of SS fractions estimated from the reference CD spectra by different methods relative to SS fractions computed from the respective reference structure. As in Fig. 3, two RDSs have been used: IDP8 for disordered proteins (blue), and SP175 for folded globular proteins (orange). The tested SS fraction estimators are: K2D3, BESTSEL, and SESCA_Bayes with four different basis sets (DS-dTSC3, DSSP-1SC3, HBSS-3SC1 and DS5-4SC1).

For the four SESCA_bayes variants using different basis sets, the smallest average RMSD^SS^ is obtained for the DS5-4SC1 basis set (6 SS classes), with 0.07 RMSD^SS^ for both globular and disordered proteins (SD of 0.04 and 0.06, respectively). The largest average RMSD^SS^ for SESCA_bayes are seen for the basis set DSSP-1SC3 (four classes), amounting to an average RMSD^SS^ of 0.12 (SD 0.06) and 0.14 (SD 0.04) for globular and disordered RDSs, respectively.

The program K2D3 estimates a three-class SS composition using a neural network that was trained on DichroCalc predictions of globular CD spectra based on their structures. K2D3 estimates globular protein SS fractions with an average RMSD^SS^ of 0.09 (SD 0.05), similar to the RMSD^SS^ SESCA_bayes achieved using the DS-dTSC3 basis set with a similar 3-class SS composition. The RMSD^SS^ of K2D3 for IDPs is 0.12 (SD 0.05), somewhat larger than that for the globular RDS. We note that the obtained SS estimation errors of K2D3 are typically small for IDPs, despite the fact that the program provides very poor back-calculated CD spectra and warns the user about the potential unreliability of those SS estimates.

The BeStSel web application provides a detailed SS estimation based on eight SS classes, four of which are associated with different types of β-sheets. An average RMSD^SS^ of 0.08 (SD 0.03) is obtained for globular proteins, and 0.14 (SD 0.05) for IDPs, which is the largest difference amongst the tested SS estimators. We attribute this difference mainly to an observed systematic overestimation of the right-handed antiparallel β-sheet (Anti3) fractions in our model IDPs (Table S9). Indeed, for the globular RDS, the SS fractions are fairly similar for BeStSel estimates and the fractions of the reference (crystal) structures. In contrast, almost none of the IDP ensemble models contains residues classified as the Anti3 class, for which BeStSel estimates fractions between 0.2 and 0.3. The only protein in the IDP8 RDS for which the Anti3 fraction was not over-estimated was *cbpn*. However, *cbpn* is a molten-globule type IDP with a stable α-helical structure, and thus its CD spectrum is more similar to those of helical proteins.

It is worth noting that BeStSel also provides a simple ordered/disordered classification of proteins based on their CD spectrum, which our IDP8 RDS also enabled us to assess. Indeed, seven of the eight proteins are correctly classified as disordered, with *cbpn* being classified as ordered. The latter result is not a true misclassification because *cbpn* is a helical molten globule and its disorder is apparent mostly on the tertiary structure level.

Further, we analyzed the SS estimates for one of the globular reference proteins, Subtilisin Carlsberg (SP175/67). The measured CD spectrum of this protein (PCDDB: CD0000067) has recently been re-measured^51^, likely because the original spectrum suffered from a severe intensity normalization error^11^. SS estimates from the original spectrum show that SESCA_bayes and BestSel both predict the protein structure rather accurately with RMSD^SS^ values between 0.02 and 0.1. In contrast, the SS estimate of K2D3 is poor with an RMSD^SS^ of 0.31 due to an overestimation of the α-helix content. For the updated CD spectrum, all three algorithms estimate accurate SS fractions with RMSD^SS^ values from 0.08 to 0.12. These results show both SESCA_bayes and BetSel are rather insensitive to normalization errors due to intensity scaling applied during their SS estimation process. In contrast, K2D3 relies on an accurate spectrum intensity, which explains inaccurate SS estimate for the original spectrum.

In contrast to the other available SS estimators, the Bayesian SS estimation method of SESCA additionally provides uncertainties for the estimated SS fractions. To test if these Bayesian uncertainties are realistic, we expressed the observed deviations to the reference SS fractions in units of *χ*^2^ analogously to *eq.* 2, but without a scaling factor. Similar to the RMSD^SS^ values above, the computed *χ*^2^ deviations also vary with the choice of the basis set. For the four basis sets, SESCA_bayes achieves average *χ*^2^ deviations for the IDP8 set of 0.87 (HBSS-3SC1), 1.03 (DS5-4SC1), 2.15 (DS-dTSC3), and 2.51 (DSSP-1SC3). Obviously, these deviations are largely within one or two Bayes standard deviations, such that the estimated uncertainty can be considered rather accurate. In contrast, the average *χ*^2^ values for the globular SP175 RDS are 1.32 (DS5-4SC3), 2.62 (HBSS-3SC1), 3.04 (DSSP-1SC3), and 5.59 (DS-dTSC3), significantly larger than for the IDP set. As the RMSD^SS^ values for the SP175 are not considerably larger than those of our IDP8 RDS, the significantly larger *χ*^2^ deviations indicate that uncertainties of the SS fractions are underestimated for the DS-DTSC3 basis set, and to a lesser extent for the DSSP-1SC3 basis set as well.

Overall, the observed RMSD^SS^ values indicate that SESCA basis sets estimate the SS composition of IDPs with a similar accuracy as globular ones, whereas the average deviation of K2D3 and BeStSel SS estimates are somewhat smaller for globular proteins and larger for IDPs. Our results also suggest that SESCA basis sets DS5-4SC1 and Ds-dTSC3 are slightly more accurate for SS estimations than HBSS-3SC1 and DSSP-1SC3, but the uncertainties of DS-dTSC3 may be underestimated.

## Conclusions

### Current method accuracy

We introduced a new reference data set (RDS) for disordered proteins comprising CD spectra of eight proteins and 14 ensemble models. This RDS, referred to as IDP8, served here to assess existing CD-based biophysical analysis methods and can also support their further development. We first determined the accuracy of the CD prediction methods SESCA, DichroCalc, and PDB2CD and compared it to their accuracy for folded globular proteins using the curated RDS SP175. Overall, the accuracy of these methods was lower (between 2.0 and 9.0 kMRE) for IDPs than for globular proteins (between 1.6 to 4.8 kMRE). SESCA predicted the CD spectra of globular and disordered proteins with a similar high accuracy; PDB2CD performed well on globular proteins but was less accurate for IDPs, whereas larger errors were seen for DichroCalc for both folded as well as disordered proteins.

Second, we used the IDP8 data set to assess the accuracy of the CD-based secondary structure estimators SESCA_bayes, K2D3, and BESTSEL. Here, the (absolute) error of SS fraction estimates was found between 0.07 and 0.14 for disordered proteins and between 0.07 and 0.12 for globular proteins. Again, the accuracy of SESCA SS estimates was similar for folded and disordered proteins. However, and in contrast to the above-mentioned CD spectrum predictions, it varied depending on the used basis set. Both K2D3 and BESTSEL provided more accurate SS estimates for globular than for disordered proteins.

Importantly, the IDP8 data set also enabled us to test if SESCA_bayes provides realistic uncertainty estimates. For the disordered proteins, the uncertainty estimates largely agreed with the actual deviations from the SS of the reference ensembles, whereas for the folded proteins, the uncertainty estimates, particularly for the smaller basis sets, tended to be smaller than the actual errors. None of the other SS estimators provides uncertainty estimates.

Over the past years, several methods for the structural characterization of folded proteins by CD spectroscopy − such as CD spectrum predictors or SS estimators – have been established and are now widely used. Their development and optimization has been enabled and driven by high quality RDSs such as SP175. Similar developments for IDPs, though pressing, have been hampered by the lack of a suitable reference data set. We addressed this obstacle by compiling IDP8, an intrinsically disordered protein RDS. Our subsequent assessments showed that the structural ensembles of IDP8 agree well with SAXS and NMR chemical shift measurements, thus establishing that their quality is sufficient for CD assessment. Using this new RDS, our assessments showed that SESCA CD predictions and SS estimations achieved similarly high accuracy for disordered proteins as we previously determined for globular proteins, which suggests that SESCA should be equally applicable to both protein classes. Further, the assessment of several other CD prediction and SS estimation methods revealed generally lower accuracy for IDPs than for globular proteins. Further, our data indicated that most of the tested methods (including SESCA) would likely benefit from re-parametrization using the IDP8 RDS. We therefore believe that our IDP8 RDS will also drive further methodological improvements in this rapidly growing field.

## Supporting information

Supplementary Information (combined)

## Data set availability

All ensemble models and CD spectra will be made publicly available through the protein ensemble database (PED) and the protein circular dichroism database (PCDDB), respectively. Until then, the CD spectra and ensemble models of the IDP8 RDS are available on request. Supplementary information about computational tool availability, precited CD spectra and estimated SS fractions are available online free of charge.

## Acknowledgements

Gabor Nagy would like to thank the Alexander von Humboldt foundation for financial support. The authors would like to thank Sonia Longhi for providing support and experimental data on *mevn*. We are thankful to Martha Brennich for the aid in SAXS measurements. We would like to thank Stefan Becker, Christian Griesinger, and Karin Müller for their help in sample preparation. We would like to thank Sarah Rauscher and Reinhardt Klement for providing simulation trajectories, experimental data, and useful discussions regarding *rsp8* and *asyn*, respectively, and Vytautas Gapsys and Tamás Lázár for helpful discussions regarding ensemble refinement and model deposition.

