## Supplementary Information (combined) for "A Reference Data Set for Circular Dichroism Spectroscopy Comprised of Validated Intrinsically Disordered Protein Models"

### Supplementary Figures:

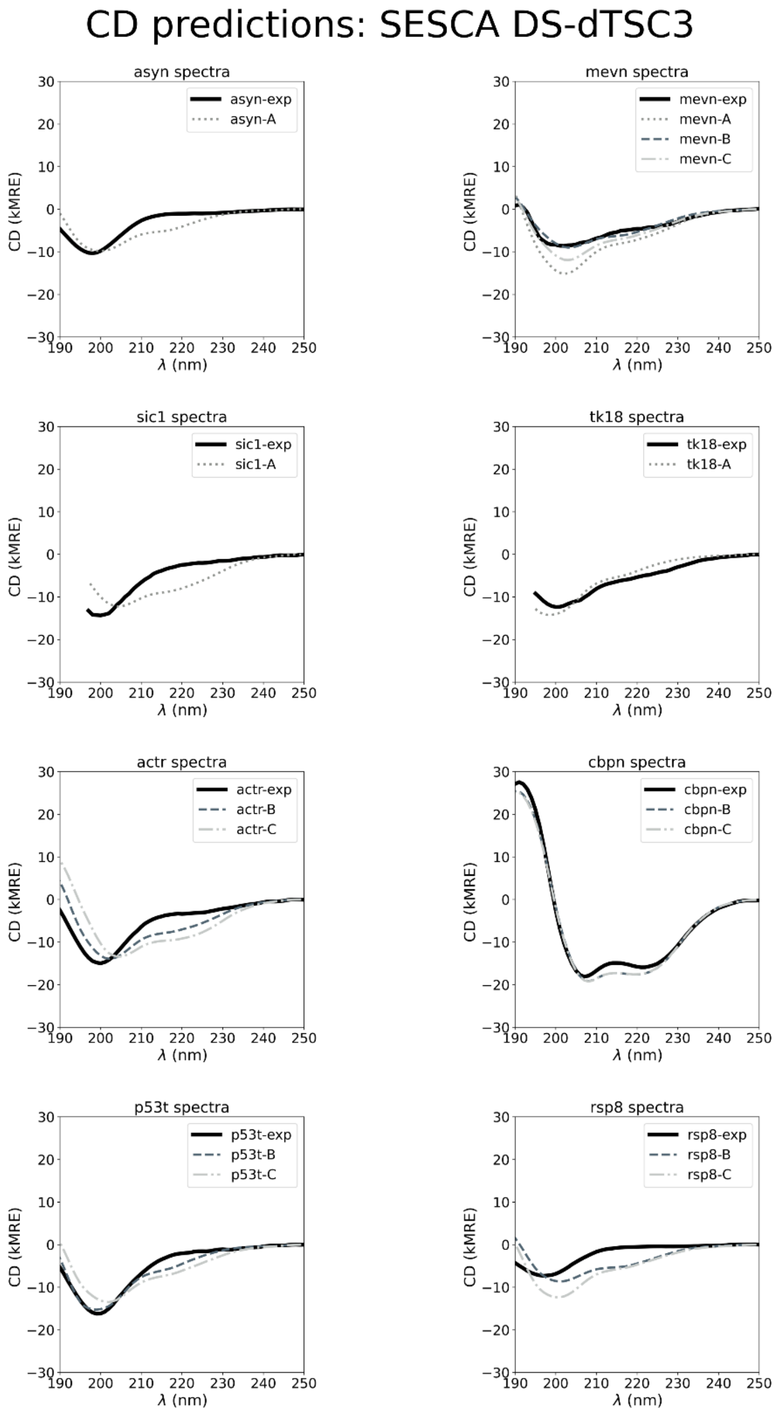


**Figure S1:** Comparison between measured CD spectra (exp), and CD spectra predicted from IDP8 ensemble models of groups A, B, and C, using the SESCA basis set DS-dTSC3.

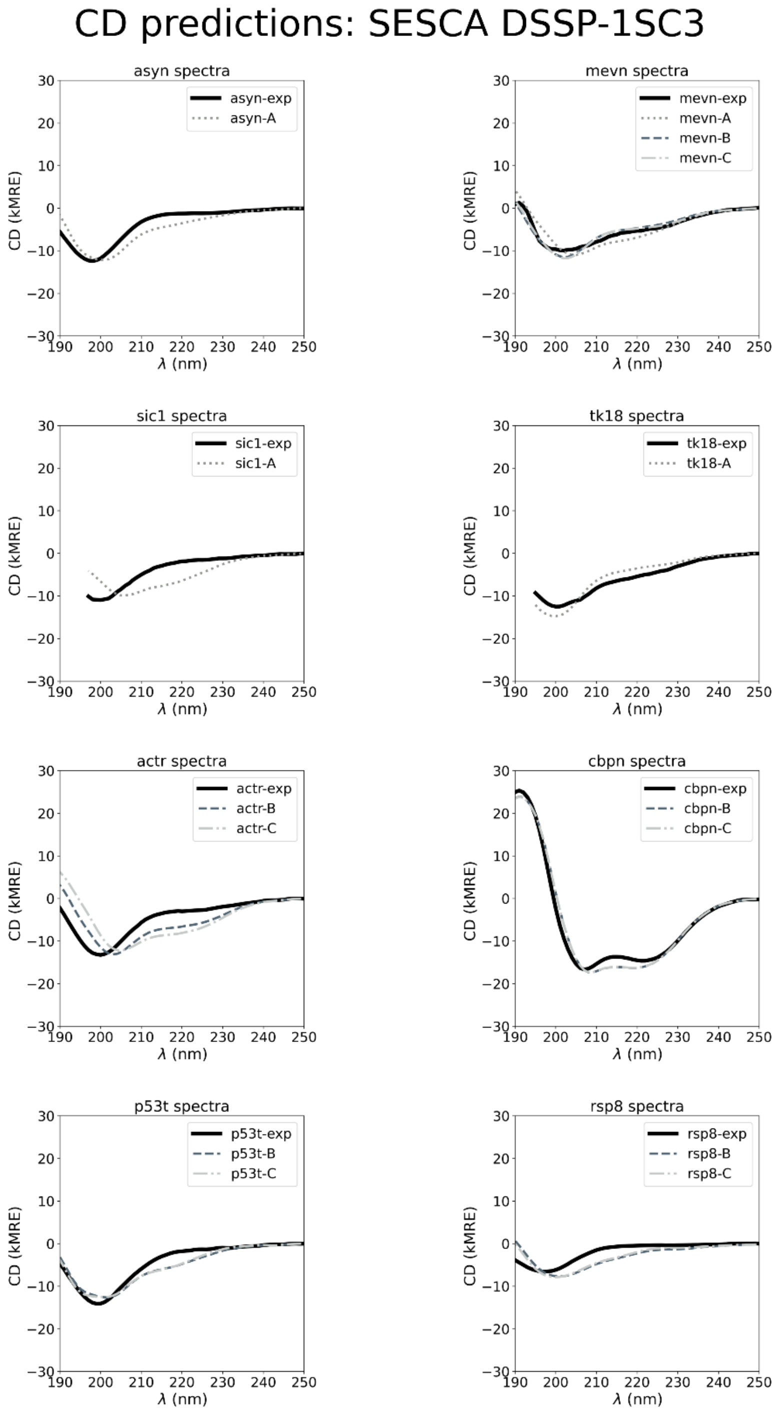


**Figure S2:** Comparison between measured CD spectra (exp), and CD spectra predicted from IDP8 ensemble models of groups A, B, and C, using the SESCA basis set DSSP-1SC3.

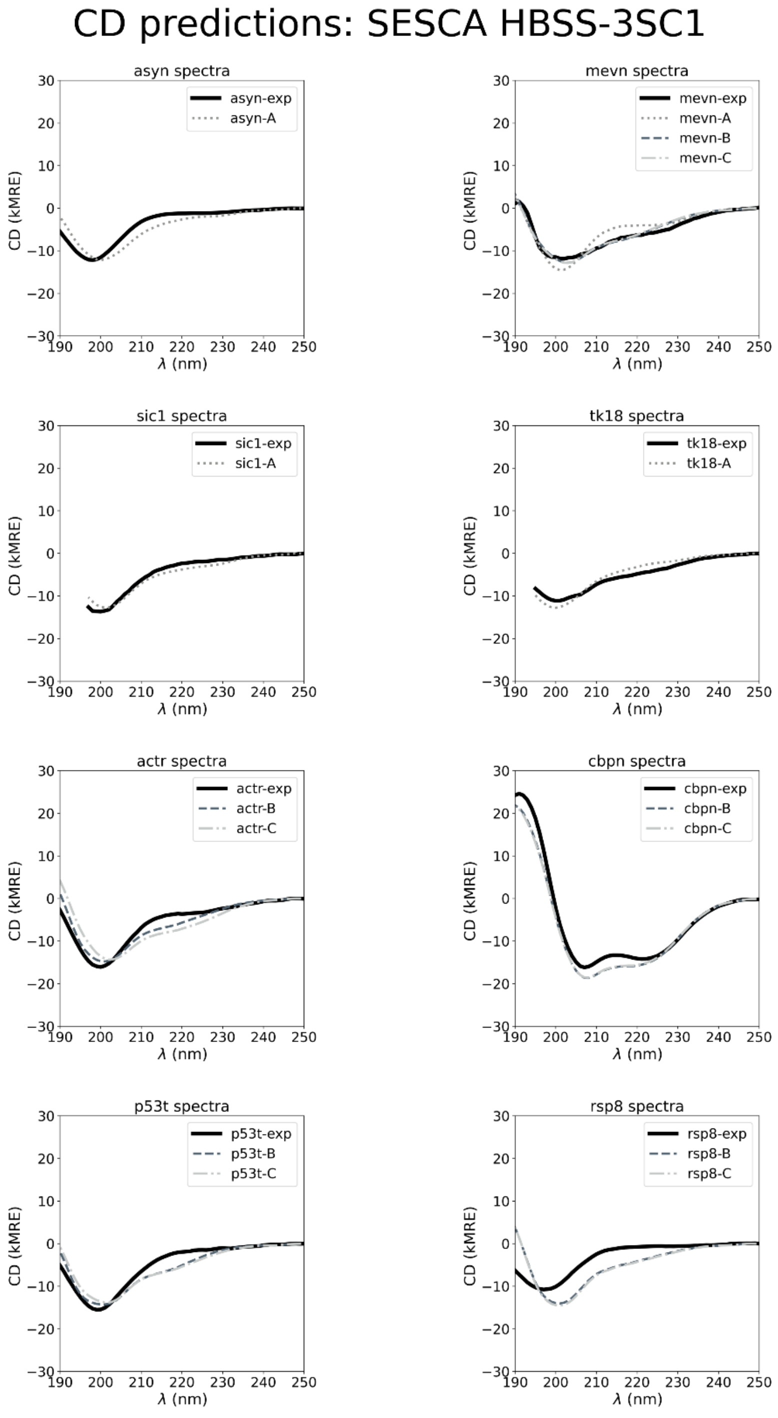


**Figure S3:** Comparison between measured CD spectra (exp), and CD spectra predicted from IDP8 ensemble models of groups A, B, and C, using the SESCA basis set HBSS-3SC1.

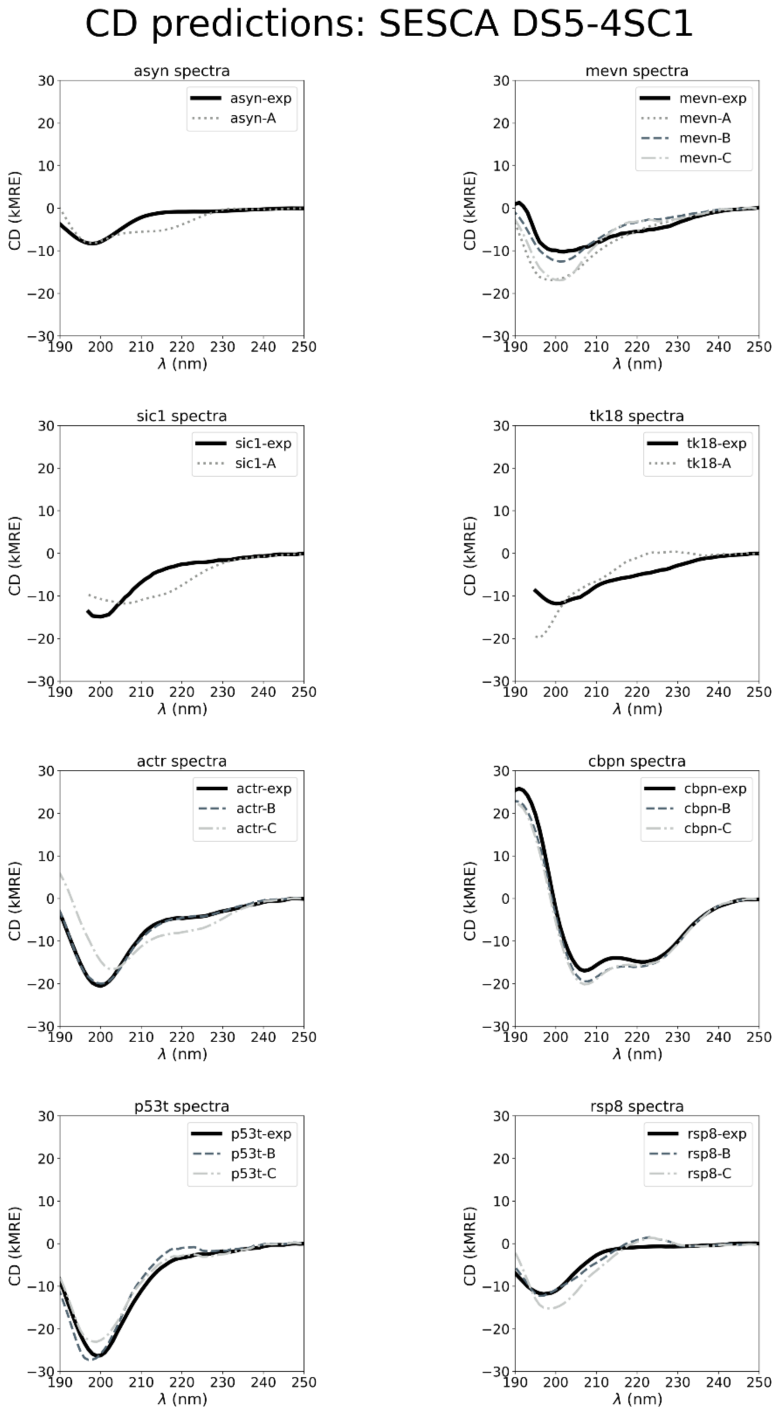


**Figure S4:** Comparison between measured CD spectra (exp), and CD spectra predicted from IDP8 ensemble models of groups A, B, and C, using the SESCA basis set DS5-4SC1.

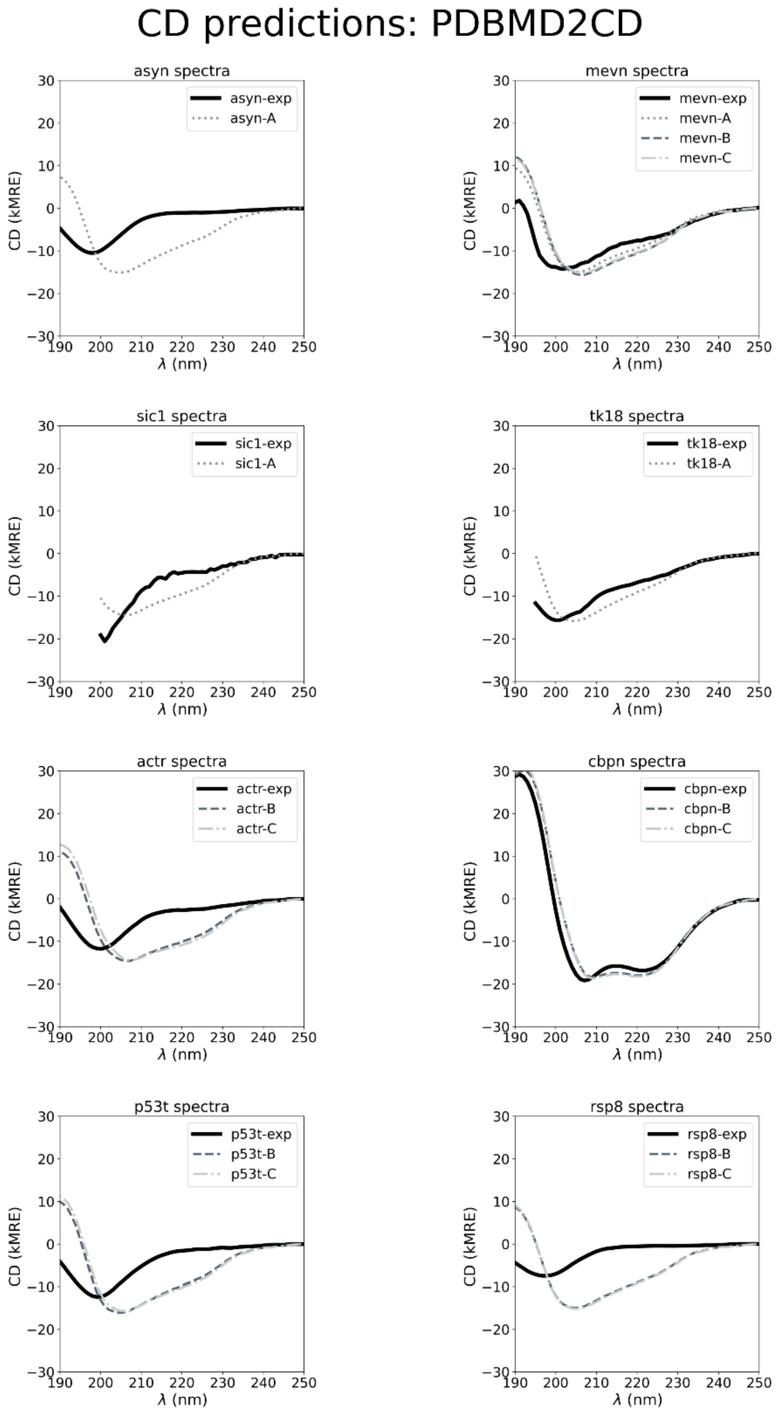


**Figure S5:** Comparison between measured CD spectra (exp), and CD spectra predicted from IDP8 ensemble models of groups A, B, and C, using the PDBMD2CD prediction server.

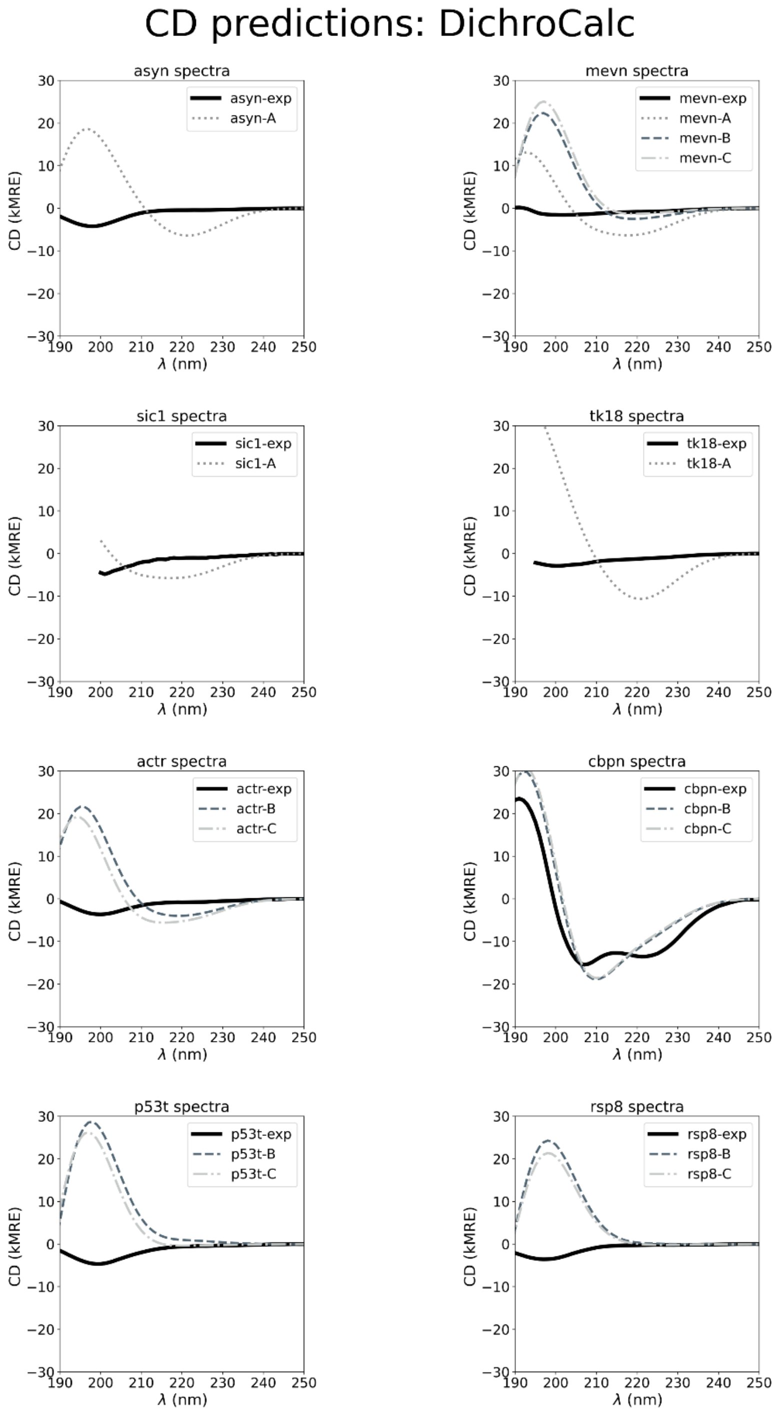


**Figure S6:** Comparison between measured CD spectra (exp), and CD spectra predicted from IDP8 ensemble models of groups A, B, and C, using the DichroCalc algorithm.

### Supplementary Tables:

**Table S1:** Bioinformatics resources. A list of online tools used in this work, including the name of the tool (resource), its main function , relevant citation (ref) used in the main publication, and URL link to access the tool.

| resource | function | ref | URL |
|---|---|---|---|
| SESCA | CD prediction | 11 | https://www.mpinat.mpg.de/sesca |
| DichroCalc | CD prediction | 5 | https://comp.chem.nottingham.ac.uk/dichrocalc |
| PDB2CD | CD prediction | 8 | https://pdb2cd.cryst.bbk.ac.uk |
| SESCA_bayes | SS estimation | 75 | https://www.mpinat.mpg.de/sesca |
| K2D3 | SS estimation | 6 | http://cbdm-01.zdv.uni-mainz.de/~andrade/k2d3 |
| BESTSEL | SS estimation | 7 | https://bestsel.elte.hu/index.php |
| SELCON3 | SS estimation | 9 | https://sites.google.com/view/sreerama |
| CCA | SS estimation | 10 | https://www.chem.elte.hu/departments/jimre |
| BRMB | database | 41 | https://bmrb.io |
| Dichroweb | database | 2 | http://dichroweb.cryst.bbk.ac.uk/html/home.shtml |
| PCDDB | database | 12 | https://pcddb.cryst.bbk.ac.uk |
| PED | database | 40 | https://proteinensemble.org |

**Table S2: MD simulations and parameters.** Summary of all MD simulations used for ensemble refinement of the newly derived IDP8 models. Different columns indicate simulation parameters for different IDP domains (abbreviations shown in the first row). The subsequent rows indicate simulation parameters used for all trajectories, temperature (T) in degrees Celsius, pressure (P) in atmospheres, and ion concentration in mol/dm$^3$ ($C_{ion}$). The rows further below describe MD trajectories, separated by horizontal lines, indicating the used force field (FF), the number of calculated trajectories (Ntraj), total simulation time (tsim), and the number of frames used for ensemble refinement (Nfr). Force fields are abbreviated as A99SB-disp (Amber99SB with dispersion corrections)[1], A99SB-ws (Amber99SB with rescaled TIP4P water)[2], A03-ws (Amber03 with rescaled water interactions)[3], A14SB-OPC (Amber14SB[4] with an OPC water model[5]), C22S-TIPS3P (CHARMM22 star with modified TIP3P water) [6], and C36M-OPC (CHARMM36M with OPC water model) [7].

| System | mevn | actr | cbpn | p53t | rsp8 |
|---|---|---|---|---|---|
| T (C°) | 25 | 25 | 25 | 25 | 25 |
| P (atm) | 1 | 1 | 1 | 1 | 1 |
| Cion (M) | 150 | 50 | 50 | 150 | 50 |
| FF | A99SB-disp | A03-ws | A03-ws | A99SB-ws | A03-ws |
| Ntraj | 3 | 2 | 2 | 30 | 1 |
| tsim (us) | 30 | 9 | 20 | 600 | 10 |
| Nfr | 12 000 | 2400 | 5000 | 12000 | 2000 |
| FF | C36M-OPC | C22S-TIPS3P | A99SB-disp | C36M-OPC | C36M-OPC |
| Ntraj | 6 | 3 | 20 | 20 | 5 |
| tsim (us) | 42 | 30 | 100 | 200 | 25 |
| Nfr | 33000 | 7500 | 20000 | 20000 | 2500 |
| FF | | | C36M-OPC | | C22S-TIPS3P |
| Ntraj | | | 20 | | 1 |
| tsim (us) | | | 100 | | 1 |
| Nfr | | | 10000 | | 1000 |
| FF | | | A14SB-OPC | | |
| Ntraj | | | 20 | | |
| tsim (us) | | | 100 | | |
| Nfr | | | 10000 | | |

**Table S3: Bayesian Maximum Entropy refinement parameters.** The table summarizes details of Bayesian maximum entropy (BME) refinement of IDP8 ensemble models. Columns denote different disordered models (abbreviations of the model shown in the first row). The subsequent rows show initial ensemble size (N0), Theta scaling parameter (Θ) and final ensemble size (Nf) for group B and group C refinements. Both sets of refinements started from the same respective initial ensemble (described in Methods) using different experimental data.

| system | mevn | actr | cbpn | p53t | rsp8 |
|---|---|---|---|---|---|
| N0 | 45 000 | 10000 | 45000 | 32000 | 5500 |
| Θ-B | 10 | 5 | 5 | 5 | 20 |
| Nf-B | 5x20 | 5x20 | 5x20 | 1X50 | 5x50 |
| Θ-C | 20 | 10 | 10 | 5 | 10 |
| Nf-C | 5X20 | 5X20 | 5x20 | 5x50 | 5x50 |

**Table S4:** Comparison between reference SS fractions and SS fractions estimated from measured CD spectra using the SESCA basis DS-dTSC3. Solid horizontal lines separate the three ensemble model groups (A, B, and C) the reference structure belongs to. Comparisons used to compute average SS estimation accuracy are shown in bold.

| DS-dTSC3 ref. model | Ref Alpha | Est Alpha | Ref Beta | Est Beta | Ref Coil | Est Coil |
|---|---|---|---|---|---|---|
| **asyn-A** | **0.02** | **0.00 (0.03)** | **0.13** | **0.03 (0.08)** | **0.85** | **0.97 (0.09)** |
| mevn-A | 0.08 | 0.03 (0.04) | 0.02 | 0.14 (0.06) | 0.90 | 0.83 (0.07) |
| **sic1-A** | **0.12** | **0.01 (0.03)** | **0.05** | **0.03 (0.07)** | **0.83** | **0.96 (0.09)** |
| **tk18-A** | **0.01** | **0.03 (0.05)** | **0.03** | **0.02 (0.07)** | **0.97** | **0.95 (0.09)** |
| **mevn-B** | **0.06** | **0.03 (0.04)** | **0.13** | **0.14 (0.06)** | **0.80** | **0.83 (0.07)** |
| **actr-B** | **0.08** | **0.01 (0.04)** | **0.05** | **0.01 (0.06)** | **0.87** | **0.98 (0.09)** |
| **cbpn-B** | **0.41** | **0.36 (0.05)** | **0.03** | **0.03 (0.05)** | **0.56** | **0.62 (0.05)** |
| **p53t-B** | **0.00** | **0.01 (0.04)** | **0.06** | **0.02 (0.07)** | **0.94** | **0.98 (0.09)** |
| **rsp8-B** | **0.03** | **0.01 (0.04)** | **0.17** | **0.06 (0.06)** | **0.80** | **0.94 (0.08)** |
| mevn-C | 0.07 | 0.03 (0.04) | 0.05 | 0.14 (0.06) | 0.87 | 0.83 (0.07) |
| actr-C | 0.15 | 0.01 (0.04) | 0.06 | 0.01 (0.06) | 0.79 | 0.98 (0.09) |
| cbpn-C | 0.41 | 0.36 (0.05) | 0.02 | 0.03 (0.05) | 0.57 | 0.62 (0.05) |
| p535t-C | 0.05 | 0.01 (0.04) | 0.06 | 0.02 (0.07) | 0.89 | 0.98 (0.09) |
| rsp8-C | 0.03 | 0.01 (0.04) | 0.08 | 0.06 (0.06) | 0.90 | 0.94 (0.08) |

**Table S5:** Comparison between reference SS fractions and SS fractions estimated from measured CD spectra using the SESCA basis DSSP-1SC3. Solid horizontal lines separate the three ensemble model groups (A, B, and C) the reference structure belongs to. Comparisons used to compute average SS estimation accuracy are shown in bold.

| DSSP-1SC3 ref. model | Ref Helix | Est Helix | Ref Strand | Est Strand | Ref Bridge | Est Bridge | Ref Other | Est Other |
|---|---|---|---|---|---|---|---|---|
| **asyn-A** | **0.01** | **0.01 (0.04)** | **0.01** | **0.13 (0.11)** | **0.01** | **0.12 (0.04)** | **0.96** | **0.74 (0.16)** |
| mevn-A | 0.11 | 0.07 (0.05) | 0.00 | 0.28 (0.11) | 0.00 | 0.07 (0.06) | 0.89 | 0.58 (0.20) |
| **sic1-A** | **0.10** | **0.06 (0.06)** | **0.00** | **0.15 (0.12)** | **0.00** | **0.09 (0.05)** | **0.89** | **0.70 (0.20)** |
| **tk18-A** | **0.00** | **0.09 (0.07)** | **0.02** | **0.09 (0.11)** | **0.02** | **0.07 (0.05)** | **0.95** | **0.76 (0.20)** |
| **mevn-B** | **0.07** | **0.07 (0.05)** | **0.02** | **0.28 (0.11)** | **0.02** | **0.07 (0.06)** | **0.90** | **0.58 (0.20)** |
| **actr-B** | **0.11** | **0.06 (0.05)** | **0.03** | **0.09 (0.10)** | **0.02** | **0.10 (0.05)** | **0.84** | **0.75 (0.17)** |
| **cbpn-B** | **0.43** | **0.46 (0.08)** | **0.00** | **0.12 (0.10)** | **0.00** | **0.08 (0.05)** | **0.57** | **0.34 (0.16)** |
| **p53t-B** | **0.02** | **0.02 (0.04)** | **0.01** | **0.08 (0.11)** | **0.01** | **0.11 (0.04)** | **0.96** | **0.79 (0.17)** |
| **rsp8-B** | **0.04** | **0.01 (0.03)** | **0.04** | **0.20 (0.11)** | **0.01** | **0.12 (0.04)** | **0.91** | **0.68 (0.15)** |
| mevn-C | 0.08 | 0.07 (0.05) | 0.02 | 0.28 (0.11) | 0.03 | 0.07 (0.06) | 0.88 | 0.58 (0.20) |
| actr-C | 0.15 | 0.06 (0.05) | 0.05 | 0.09 (0.10) | 0.01 | 0.10 (0.05) | 0.79 | 0.75 (0.17) |
| cbpn-C | 0.43 | 0.46 (0.08) | 0.00 | 0.12 (0.10) | 0.00 | 0.08 (0.05) | 0.57 | 0.34 (0.16) |
| p535t-C | 0.00 | 0.02 (0.04) | 0.00 | 0.08 (0.11) | 0.00 | 0.11 (0.04) | 1.00 | 0.79 (0.17) |
| rsp8-C | 0.03 | 0.01 (0.03) | 0.04 | 0.20 (0.11) | 0.01 | 0.12 (0.04) | 0.92 | 0.68 (0.15) |

**Table S6:** Comparison between reference SS fractions and SS fractions estimated from measured CD spectra using the SESCA basis HBSS-3SC1. Solid horizontal lines separate the three ensemble model groups (A, B, and C) the reference structure belongs to. Comparisons used to compute average SS estimation accuracy are shown in bold.

| HBSS-3SC1 ref. model | Ref Helix_reg | Est Helix_reg | Ref Helix_irr | Est Helix_irr | Ref Beta_all | Est Beta_all | Ref Turn | Est Turn | Ref Other | Est Other |
|---|---|---|---|---|---|---|---|---|---|---|
| **asyn-A** | **0.00** | **0.00 (0.03)** | **0.00** | **0.02 (0.04)** | **0.00** | **0.02 (0.06)** | **0.49** | **0.46 (0.29)** | **0.50** | **0.50 (0.30)** |
| mevn-A | 0.02 | 0.01 (0.03) | 0.05 | 0.08 (0.08) | 0.00 | 0.07 (0.07) | 0.57 | 0.37 (0.24) | 0.36 | 0.46 (0.22) |
| **sic1-A** | **0.02** | **0.01 (0.03)** | **0.02** | **0.04 (0.06)** | **0.00** | **0.02 (0.05)** | **0.47** | **0.33 (0.22)** | **0.49** | **0.60 (0.24)** |
| **tk18-A** | **0.00** | **0.02 (0.03)** | **0.00** | **0.05 (0.06)** | **0.01** | **0.02 (0.04)** | **0.39** | **0.21 (0.22)** | **0.60** | **0.71 (0.24)** |
| **mevn-B** | **0.06** | **0.01 (0.03)** | **0.01** | **0.08 (0.08)** | **0.02** | **0.07 (0.07)** | **0.09** | **0.37 (0.24)** | **0.83** | **0.46 (0.22)** |
| **actr-B** | **0.02** | **0.01 (0.02)** | **0.01** | **0.03 (0.05)** | **0.03** | **0.02 (0.05)** | **0.18** | **0.30 (0.20)** | **0.75** | **0.65 (0.21)** |
| **cbpn-B** | **0.35** | **0.33 (0.06)** | **0.02** | **0.09 (0.08)** | **0.00** | **0.02 (0.04)** | **0.16** | **0.27 (0.18)** | **0.46** | **0.29 (0.16)** |
| **p53t-B** | **0.00** | **0.00 (0.02)** | **0.01** | **0.02 (0.05)** | **0.01** | **0.02 (0.04)** | **0.08** | **0.45 (0.28)** | **0.91** | **0.51 (0.28)** |
| **rsp8-B** | **0.01** | **0.00 (0.02)** | **0.01** | **0.01 (0.04)** | **0.03** | **0.02 (0.05)** | **0.18** | **0.44 (0.25)** | **0.78** | **0.52 (0.26)** |
| mevn-C | 0.05 | 0.01 (0.03) | 0.01 | 0.08 (0.08) | 0.02 | 0.07 (0.07) | 0.10 | 0.37 (0.24) | 0.82 | 0.46 (0.22) |
| actr-C | 0.07 | 0.01 (0.02) | 0.02 | 0.03 (0.05) | 0.02 | 0.02 (0.05) | 0.18 | 0.30 (0.20) | 0.71 | 0.65 (0.21) |
| cbpn-C | 0.35 | 0.33 (0.06) | 0.02 | 0.09 (0.08) | 0.00 | 0.02 (0.04) | 0.16 | 0.27 (0.18) | 0.48 | 0.29 (0.16) |
| p535t-C | 0.01 | 0.00 (0.02) | 0.02 | 0.02 (0.05) | 0.01 | 0.02 (0.04) | 0.11 | 0.45 (0.28) | 0.85 | 0.51 (0.28) |
| rsp8-C | 0.01 | 0.00 (0.02) | 0.01 | 0.01 (0.04) | 0.01 | 0.02 (0.05) | 0.18 | 0.44 (0.25) | 0.79 | 0.52 (0.26) |

**Table S7:** Comparison between reference SS fractions and SS fractions estimated from measured CD spectra using the SESCA basis DS5-4SC1. Solid horizontal lines separate the three ensemble model groups (A, B, and C) the reference structure belongs to. Comparisons used to compute average SS estimation accuracy are shown in bold.

| DS5-4SC1 ref. model | Ref Helix1 | Est Helix1 | Ref Helix2 | Est Helix2 | Ref Beta1 | Est Beta1 | Ref Turn1 | Est Turn1 | Ref Turn2 | Est Turn2 | Ref Other | Est Other |
|---|---|---|---|---|---|---|---|---|---|---|---|---|
| **asyn-A** | **0.02** | **0.02 (0.03)** | **0.01** | **0.04 (0.05)** | **0.13** | **0.05 (0.06)** | **0.11** | **0.38 (0.08)** | **0.19** | **0.30 (0.11)** | **0.55** | **0.20 (0.15)** |
| mevn-A | 0.05 | 0.04 (0.05) | 0.06 | 0.07 (0.07) | 0.02 | 0.13 (0.07) | 0.25 | 0.20 (0.09) | 0.19 | 0.29 (0.12) | 0.43 | 0.27 (0.18) |
| **sic1-A** | **0.08** | **0.04 (0.04)** | **0.05** | **0.06 (0.06)** | **0.05** | **0.07 (0.07)** | **0.13** | **0.35 (0.17)** | **0.16** | **0.26 (0.18)** | **0.54** | **0.22 (0.18)** |
| **tk18-A** | **0.01** | **0.09 (0.06)** | **0.00** | **0.07 (0.07)** | **0.03** | **0.05 (0.06)** | **0.31** | **0.36 (0.14)** | **0.09** | **0.19 (0.13)** | **0.57** | **0.24 (0.19)** |
| **mevn-B** | **0.05** | **0.04 (0.05)** | **0.02** | **0.07 (0.07)** | **0.13** | **0.13 (0.07)** | **0.23** | **0.20 (0.09)** | **0.29** | **0.29 (0.12)** | **0.27** | **0.27 (0.18)** |
| **actr-B** | **0.05** | **0.07 (0.05)** | **0.05** | **0.03 (0.04)** | **0.05** | **0.03 (0.05)** | **0.27** | **0.31 (0.10)** | **0.29** | **0.30 (0.11)** | **0.29** | **0.26 (0.16)** |
| **cbpn-B** | **0.36** | **0.35 (0.07)** | **0.06** | **0.03 (0.04)** | **0.03** | **0.04 (0.05)** | **0.15** | **0.13 (0.08)** | **0.20** | **0.25 (0.10)** | **0.19** | **0.20 (0.14)** |
| **p53t-B** | **0.00** | **0.05 (0.04)** | **0.02** | **0.02 (0.04)** | **0.06** | **0.02 (0.04)** | **0.39** | **0.37 (0.08)** | **0.32** | **0.31 (0.11)** | **0.21** | **0.22 (0.14)** |
| **rsp8-B** | **0.01** | **0.02 (0.03)** | **0.08** | **0.04 (0.05)** | **0.17** | **0.09 (0.07)** | **0.31** | **0.38 (0.08)** | **0.29** | **0.25 (0.10)** | **0.15** | **0.22 (0.15)** |
| mevn-C | 0.06 | 0.04 (0.05) | 0.03 | 0.07 (0.07) | 0.05 | 0.13 (0.07) | 0.29 | 0.20 (0.09) | 0.31 | 0.29 (0.12) | 0.26 | 0.27 (0.18) |
| actr-C | 0.12 | 0.07 (0.05) | 0.05 | 0.03 (0.04) | 0.06 | 0.03 (0.05) | 0.18 | 0.31 (0.10) | 0.29 | 0.30 (0.11) | 0.31 | 0.26 (0.16) |
| cbpn-C | 0.37 | 0.35 (0.07) | 0.05 | 0.03 (0.04) | 0.02 | 0.04 (0.05) | 0.18 | 0.13 (0.08) | 0.20 | 0.25 (0.10) | 0.18 | 0.20 (0.14) |
| p535t-C | 0.03 | 0.05 (0.04) | 0.04 | 0.02 (0.04) | 0.06 | 0.02 (0.04) | 0.35 | 0.37 (0.08) | 0.31 | 0.31 (0.11) | 0.22 | 0.22 (0.14) |
| rsp8-C | 0.02 | 0.02 (0.03) | 0.03 | 0.04 (0.05) | 0.08 | 0.09 (0.07) | 0.25 | 0.38 (0.08) | 0.32 | 0.25 (0.10) | 0.30 | 0.22 (0.15) |

**Table S8:** Comparison between reference SS fractions and SS fractions estimated from measured CD spectra using the K2D3 web application. Solid horizontal lines separate the three ensemble model groups (A, B, and C) the reference structure belongs to. Comparisons used to compute average SS estimation accuracy are shown in bold.

| K2D3 ref. model | Ref Alpha | Est Alpha | Ref Beta | Est Beta | Ref Coil | Est Coil |
|---|---|---|---|---|---|---|
| **asyn-A** | **0.02** | **0.04** | **0.13** | **0.32** | **0.85** | **0.65** |
| mevn-A | 0.08 | 0.03 | 0.02 | 0.28 | 0.90 | 0.68 |
| **sic1-A** | **0.12** | **0.06** | **0.05** | **0.21** | **0.83** | **0.73** |
| **tk18-A** | **0.01** | **0.07** | **0.03** | **0.22** | **0.97** | **0.71** |
| **mevn-B** | **0.06** | **0.03** | **0.13** | **0.28** | **0.80** | **0.68** |
| **actr-B** | **0.08** | **0.03** | **0.05** | **0.20** | **0.87** | **0.77** |
| **cbpn-B** | **0.41** | **0.46** | **0.03** | **0.10** | **0.56** | **0.44** |
| **p53t-B** | **0.00** | **0.03** | **0.06** | **0.21** | **0.94** | **0.76** |
| **rsp8-B** | **0.03** | **0.03** | **0.17** | **0.11** | **0.80** | **0.86** |
| mevn-C | 0.07 | 0.03 | 0.05 | 0.28 | 0.87 | 0.68 |
| actr-C | 0.15 | 0.03 | 0.06 | 0.20 | 0.79 | 0.77 |
| cbpn-C | 0.41 | 0.46 | 0.02 | 0.10 | 0.57 | 0.44 |
| p535t-C | 0.05 | 0.03 | 0.06 | 0.21 | 0.89 | 0.76 |
| rsp8-C | 0.03 | 0.03 | 0.08 | 0.11 | 0.90 | 0.86 |

**Table S9:** Comparison between reference SS fractions and SS fractions estimated from measured CD spectra using the BESTSEL web application. Solid horizontal lines separate the three ensemble model groups (A, B, and C) the reference structure belongs to. Comparisons used to compute average SS estimation accuracy are shown in bold.

| BESTSEL ref. model | Ref Helix | Est Helix | Ref Anti1 | Est Anti1 | Ref Anti2 | Est Anti2 | Ref Anti3 | Est Anti3 | Ref Parallel | Est Parallel | Ref Other | Est Other |
|---|---|---|---|---|---|---|---|---|---|---|---|---|
| **asyn-A** | **0.00** | **0.03** | **0.00** | **0.00** | **0.00** | **0.02** | **0.00** | **0.29** | **0.00** | **0.00** | **1.00** | **0.67** |
| mevn-A | 0.02 | 0.08 | 0.00 | 0.00 | 0.00 | 0.12 | 0.00 | 0.20 | 0.00 | 0.03 | 0.98 | 0.57 |
| **sic1-A** | **0.02** | **0.05** | **0.00** | **0.00** | **0.00** | **0.00** | **0.00** | **0.22** | **0.00** | **0.02** | **0.98** | **0.72** |
| **tk18-A** | **0.00** | **0.11** | **0.00** | **0.00** | **0.00** | **0.00** | **0.00** | **0.22** | **0.00** | **0.00** | **0.99** | **0.67** |
| **mevn-B** | **0.06** | **0.08** | **0.00** | **0.00** | **0.00** | **0.12** | **0.00** | **0.20** | **0.02** | **0.03** | **0.93** | **0.57** |
| **actr-B** | **0.02** | **0.08** | **0.00** | **0.00** | **0.00** | **0.00** | **0.01** | **0.20** | **0.02** | **0.00** | **0.95** | **0.72** |
| **cbpn-B** | **0.35** | **0.39** | **0.00** | **0.00** | **0.00** | **0.00** | **0.00** | **0.03** | **0.00** | **0.00** | **0.65** | **0.58** |
| **p53t-B** | **0.00** | **0.07** | **0.00** | **0.00** | **0.00** | **0.01** | **0.00** | **0.26** | **0.01** | **0.00** | **0.99** | **0.66** |
| **rsp8-B** | **0.01** | **0.04** | **0.00** | **0.00** | **0.00** | **0.03** | **0.01** | **0.24** | **0.02** | **0.00** | **0.97** | **0.70** |
| mevn-C | 0.05 | 0.08 | 0.00 | 0.00 | 0.00 | 0.12 | 0.00 | 0.20 | 0.02 | 0.03 | 0.93 | 0.57 |
| actr-C | 0.07 | 0.08 | 0.00 | 0.00 | 0.00 | 0.00 | 0.00 | 0.20 | 0.02 | 0.00 | 0.91 | 0.72 |
| cbpn-C | 0.35 | 0.39 | 0.00 | 0.00 | 0.00 | 0.00 | 0.00 | 0.03 | 0.00 | 0.00 | 0.65 | 0.58 |
| p535t-C | 0.01 | 0.07 | 0.00 | 0.00 | 0.00 | 0.01 | 0.00 | 0.26 | 0.01 | 0.00 | 0.98 | 0.66 |
| rsp8-C | 0.01 | 0.04 | 0.00 | 0.00 | 0.00 | 0.03 | 0.01 | 0.24 | 0.01 | 0.00 | 0.98 | 0.70 |

**Table S10:** Computed RMSD$^{SS}$ values per reference model. Root-mean-squared deviations were computed from SS fractions estimated from the measured CD spectra by a given method (columns) and the SS fractions of a corresponding refence ensemble (rows). Solid horizontal lines separate the three ensemble model groups (A, B, and C). SS fractions for all SESCA basis sets (DS-dTSC3, DSSP-1SC3 HBSS-3SC1, and DS5-4SC1) as well as the SS estimation methods K2D3 and BESTSEL are shown in Tables S2-S6. The reference models used to compute the average accuracy of each method is shown in bold.

| ref. model | DS-dTSC3 | DSSP-1SC3 | HBSS-3SC1 | DS5-4SC1 | K2D3 | BESTSEL |
|---|---|---|---|---|---|---|
| **asyn-A** | **0.09** | **0.14** | **0.02** | **0.19** | **0.16** | **0.18** |
| mevn-A | 0.08 | 0.21 | 0.11 | 0.09 | 0.20 | 0.19 |
| **sic1-A** | **0.10** | **0.13** | **0.08** | **0.17** | **0.11** | **0.14** |
| **tk18-A** | **0.02** | **0.11** | **0.10** | **0.15** | **0.19** | **0.17** |
| **mevn-B** | **0.03** | **0.21** | **0.21** | **0.03** | **0.11** | **0.17** |
| **actr-B** | **0.08** | **0.07** | **0.07** | **0.02** | **0.11** | **0.13** |
| **cbpn-B** | **0.05** | **0.14** | **0.09** | **0.03** | **0.07** | **0.03** |
| **p53t-B** | **0.03** | **0.11** | **0.24** | **0.02** | **0.14** | **0.18** |
| **rsp8-B** | **0.10** | **0.15** | **0.16** | **0.06** | **0.05** | **0.15** |
| mevn-C | 0.06 | 0.20 | 0.21 | 0.05 | 0.18 | 0.17 |
| actr-C | 0.14 | 0.07 | 0.07 | 0.06 | 0.11 | 0.11 |
| cbpn-C | 0.04 | 0.14 | 0.10 | 0.04 | 0.07 | 0.04 |
| p53t-C | 0.06 | 0.13 | 0.21 | 0.02 | 0.11 | 0.17 |
| rsp8-C | 0.03 | 0.16 | 0.17 | 0.07 | 0.03 | 0.15 |
